# Class A capsid assembly modulator apoptotic elimination of hepatocytes with high HBV core antigen level in vivo is dependent on de novo core protein translation

**DOI:** 10.1101/2023.09.25.559252

**Authors:** Jan Martin Berke, Ying Tan, Sarah Sauviller, Dai-tze Wu, Ke Zhang, Nádia Conceição-Neto, Alfonso Blázquez Moreno, Desheng Kong, George Kukolj, Chris Li, Ren Zhu, Isabel Nájera, Frederik Pauwels

## Abstract

**Background and aims:** Capsid assembly (CA) is a critical step in the hepatitis B virus (HBV) life cycle, mediated by the viral core protein. CA is the target for various new anti-viral candidate therapeutics known as capsid assembly modulators (CAMs) of which the CAM-aberrant (CAM-A) class induces aberrant shaped core protein structures and lead to hepatocyte cell death. The aim of the studies was to identify the mechanism of action of the CAM-A modulators leading to HBV infected hepatocyte elimination.

**Methods:** The CAM-A mediated mechanism of HBsAg reduction was evaluated in vitro in a stable HBV replicating cell line and in vivo in AAV-HBV transduced C57BL/6, C57BL/6 SCID and HBV-infected chimeric mice with humanized livers.

**Results:** In vivo treatment with CAM-A modulators induced pronounced reductions in HBe- and HBsAg which were associated with a transient increase in ALT. Both HBs- and HBeAg reduction and ALT increase were delayed in C57BL/6 SCID and chimeric mice, suggesting that adaptive immune responses may indirectly contribute to this phenotype. However, depletion of CD8+ T-cells in transduced wild-type mice did not have a negative impact on antigen reduction, indicating that CD8+ T-cell responses are not essential.

Coinciding with the transient ALT elevation in AAV-HBV transduced mice, we observed a transient increase in markers related to endoplasmic reticulum stress and apoptosis as well as cytokines related to apoptosis pathways, followed by the detection of a proliferation marker. Pathway enrichment analysis of microarray data revealed that antigen presentation pathway (MHC-I) was upregulated, overlapping with observed apoptosis. Combination treatment with HBV-specific siRNA demonstrated that CAM-A mediated HBsAg reduction is dependent on de novo core protein translation and that the effect is dependent on high levels of core protein expression, which will likely focus the CHB sub-population that could respond.

**Conclusion:** CAM-A treatment eradicates HBV infected hepatocytes with high core protein levels through the induction of apoptosis a promising approach as part of a regimen to achieve functional cure.

**Lay summary:** Treatment with hepatitis B virus (HBV) capsid assembly modulators that induce the formation of aberrant HBV core protein structures (CAM-A) leads to programmed cell death, apoptosis, of HBV-infected hepatocytes and subsequent reduction of HBV antigens, which differentiates CAM-A from other CAMs. The effect is dependent on the *de novo* synthesis and high levels of core protein.

## INTRODUCTION

Hepatitis B is a major global health problem, with approximately 2 billion people estimated to have had an acute infection with the hepatitis B virus (HBV). The World Health Organization (WHO) have estimated that 296 million people were living with chronic hepatitis B (CHB) infection in 2019, with 1.5 million new infections each year (1). 1–5 % of those infected as adults with HBV develop chronic infection, while the remainder clear the infection. Approximately 25–50 % of children infected with HBV at ages 1–5 develop chronic infection, as do >90 % of infants infected at birth (2). Approximately 20–30% of individuals with chronic hepatitis B (CHB) ultimately develop cirrhosis, liver failure or hepatocellular carcinoma (1). Current CHB treatments rarely result in a functional cure, defined as sustained loss of hepatitis B surface antigen (HBsAg) with or without HBsAg seroconversion and undetectable HBV DNA in serum 24 weeks off treatment (3). Finite treatment with an immunomodulator (pegylated interferon-α), can induce loss of HBV surface antigen (HBsAg) but only in a limited patient subset (4, 5) and due to side effects may be poorly tolerated (6). Treatment with a nucleos(t)ide analogue (NA; e.g. tenofovir disoproxil [TDF], or entecavir [ETV]) is generally life-long, well tolerated (6) and associated with a high barrier to resistance. Nucleos(t)ide treatment controls viral replication, eliminates liver inflammation and prevents cirrhosis, but only partially reduces the risk of hepatocellular carcinoma (7, 8, 9, 10), and rarely leads to functional cure.

Thus, novel therapeutic approaches with improved efficacy and an acceptable side effect profile are required to provide finite treatment regimens that can achieve high functional cure rates in CHB.

HBV capsid assembly is a critical step in the HBV’s life cycle and hence represents a therapeutic target. One of the seven proteins encoded by HBV is core, a protein which self-assembles to form the viral nucleocapsid. Core plays a role in most steps of the HBV life cycle e.g. subcellular trafficking and transport, release of the HBV genome, capsid assembly, and reverse transcription (11). The HBV life cycle has been well described (11). In brief, the first cytoplasmic step is formation of a capsid containing pre-genomic RNA (pgRNA) and viral polymerase, initiated by polymerase-bound pgRNA (pol-pgRNA) association with three core protein dimers (nucleation). Single dimers are rapidly added to this nucleus to form the nucleic acid containing capsid or nucleocapsid (12). Within the nucleocapsid, reverse transcription of the pgRNA by viral polymerase produces relaxed circular DNA (rcDNA). The rcDNA-containing capsid either shuttles to the nucleus to replenish cccDNA or becomes enveloped, thereby forming an infectious viral particle to be released from the cell (13).

Capsid assembly modulators (CAMs) accelerate the kinetics of capsid assembly, whereby they prevent pol-pgRNA complex encapsidation and block HBV replication (14, 15). CAMs also interfere with cccDNA transcription/de-novo formation during the early steps of infection (16, 17, 18). Such characteristics differentiate CAMs from NAs, as the latter solely interfere with the reverse transcription and polymerase process, inhibiting HBV replication. There are several types of CAMs which impact pol-pgRNA encapsidation at the nucleation step and these can be broadly categorized into two classes. CAMs whose MOA results in the formation of empty, morphologically intact capsids are referred to as CAM-E (E = empty capsids) and CAMs whose MOA results in the formation of pleiomorphic non-capsid structures (i.e. aberrant particles) are referred to as CAM-A (A = aberrant structure) (19). Examples of the latter class include heteroaryldihydropyrimidine (HAP) chemotypes such as BAY41-4109 (20, 21), GLS4 (15), JNJ-890 (18) and RO7049389 (22, 23). Interestingly, treatment of AAV-HBV transduced mice with RO7049389 not only inhibited HBV DNA replication, but also caused a strong decrease of HBe- and HBsAg (22). Several CAM-Es and CAM-As have entered clinical trials, but meaningful HBsAg reductions have not been observed (for an overview see (24)).

In this paper, we studied the underlying biology of CAM-A modulators induced HBsAg loss in vitro and in vivo in AAV-HBV-transduced and HBV-infected chimeric mice with humanized livers. The in vivo dosage for each CAM-A modulator was determined based on public information and previous pharmacokinetic studies.

## MATERIALS AND METHODS

Please see Supplementary information for detailed materials and methods.

## RESULTS

### HAP-1, 2, 3 have anti-HBV activity without inducing cytotoxicity *in vitro*

Three different HAP CAM-A compounds (HAP-1, HAP-2, and HAP-3) with favorable pharmacokinetic and ADME (absorption, distribution, metabolism, excretion) profiles were used to study the underlying biology of CAM-A induced HBsAg reduction. The anti-HBV activity of HAP-1, 2 and 3 was evaluated in stable core inducible HBV-replicating HepG2.117 cells that can be induced for HBV replication (25). The mean 50% effective concentrations (EC_50_) of HAP-1, 2 and 3 were 86, 151 and 35 nM, respectively (Table 1). HAP-1, 2 and 3 did not show cytotoxicity up to 50 µM, the highest concentration tested, in HepG2 cells (selectivity index >331; Table 1).

**Table 1.**
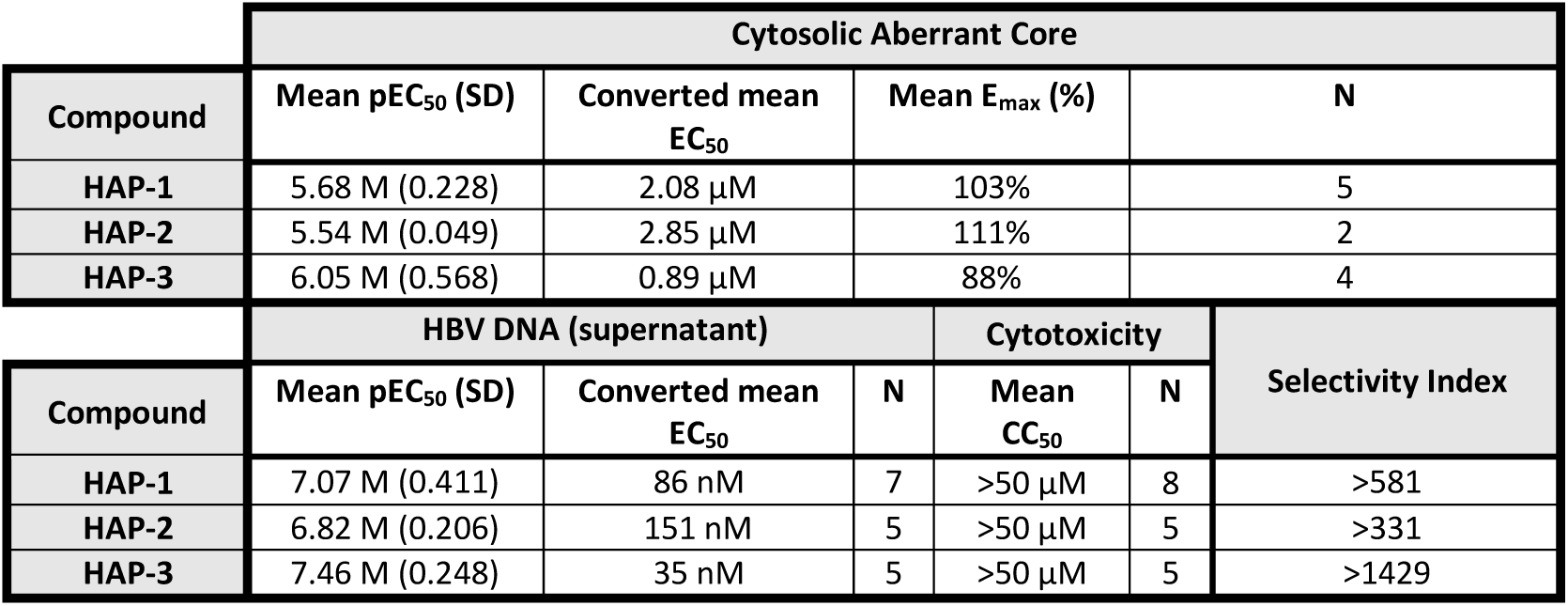
Antiviral properties and cytotoxicity of HAP-1, HAP-2 and HAP-3 in HepG2.117. . The antiviral properties of HAP-1, 2 and 3 were tested in a dose-response assay in stable HBV-replicating HepG2.117 cells. The potency of HAP-1, 2 and 3 to induce aberrant cytosolic core protein structures was assessed in a high content imaging assay with the EC_50_ representing the 50% effective concentration and the E_max_ effect as the maximum effect induced compared to a HAP reference compound (GLS-4, 1 µM). Intracellular HBV DNA was extracted from cell lysates and assessed using qPCR. The cytotoxicity was assessed in HepG2 cells using an ATP-lite read-out. In each experiment, EC_50_ values were determined based on the mean core induction/inhibition from two wells per compound concentration. pEC_50_: negative logarithm of the EC_50_; EC_50_: 50% effective concentration; SD: Standard deviation; CC_50_: 50% cytotoxic concentration; E_max_: maximal effect to induce core aberrant structures; N: number of experiments; qPCR: quantitative polymerase chain reaction; SI: selectivity index (= CC_50_ /HBV DNA EC_50_).

To confirm that HAP-1, 2 and 3 induced aberrant HBV core protein the compounds were tested in a high-content imaging assay in HepG2.117 cells designed to identify and quantify the induction of aberrant HBV core proteins (26). HAP-1, 2 and 3 induced aberrant cytosolic core protein (represented by “dot-like structures”) formation with a mean 50% effective concentrations of 2.08, 2.85 and 0.89 µM, respectively (Table 1), confirming that the compounds possessed a CAM-A MOA (Figure 1A-E).

**Figure 1.**
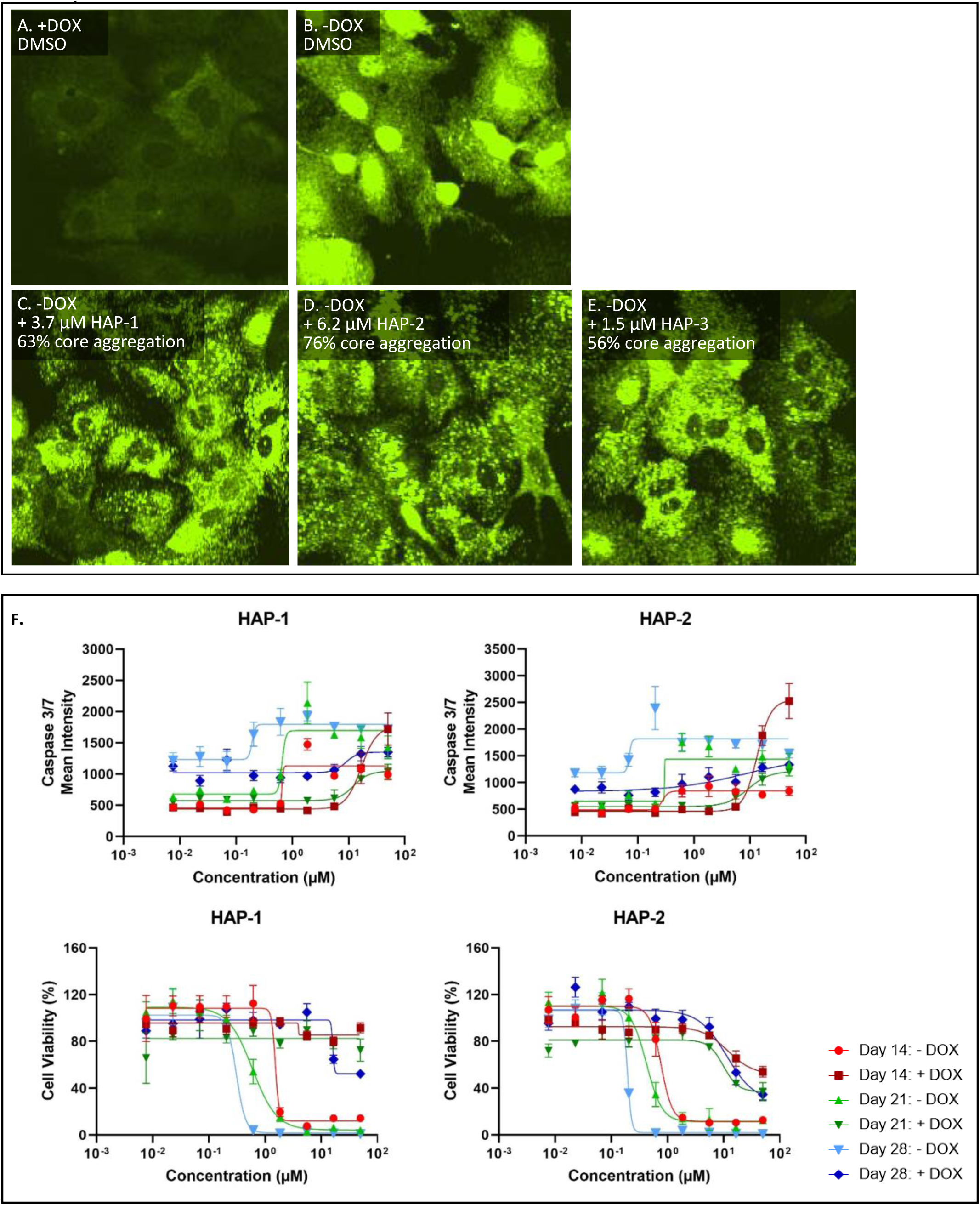
Formation of aberrant HBV core protein structures induced by HAP-1, HAP-2 and HAP-3. HepG2.117 cells were not induced (**A**, +DOX) or induced **(B-E**, -DOX) for core expression or treated with DMSO as a control **(A, B)**, HAP-1 **(C)**, HAP-2 **(D)** or HAP-3 **(E)**. Core protein was immunolabeled with primary core antibody and fluorescently labeled with Alexa 488 secondary antibody. Fluorescence was detected by confocal imaging. **(B)** A uniform core distribution in the nucleus and cytoplasm was observed in HepG2.117 cells induced for core expression (-DOX) and treated with DMSO. **(C, D, E)** Treatment of HepG2.117 with HAP-1, 2 and 3 led to the formation of aberrant core structures in the cytoplasm (represented by “dot-like structures”). The formation of aberrant cytosolic core structures was normalized to a reference HAP compound (GLS-4 used at a concentration of 1 µM) as 100% effect and DMSO treatment as 0% effect. The representative images show the induced effect around the EC_50_ concentration to induce aberrant core structures. **(F)** Induction of Caspase 3/7 activity and cell viability evaluation of HepG2.117 cells that were not induced (+DOX) or induced (-DOX) for core expression and treated with HAP-1 or HAP-2 for 28 days. Caspase activity and cell viability was assessed on day 14, day 21 and day 28. In each graph data are shown as mean ± SEM.

The formation of intracellular aberrant core protein structures could lead to cellular stress responses and, if aberrant core accumulates, to apoptosis. To test if CAM-A-mediated aberrant core protein formation induced apoptosis, HepG2.117 cells were cultured with (suppressed HBV replication) or without (induced HBV replication) doxycycline in presence of HAP-1 and 2 for up to 4 weeks and a caspase 3/7 assay was performed to detect apoptotic cells. A dose-dependent caspase 3/7 activity was only detected in HBV replicating, but not in HBV replication-suppressed HepG2.117 cells, after 2 weeks of treatment and progressed further at 3 and 4 weeks of treatment. The increase in caspase 3/7 activity was associated with a decrease in cell viability, which was evaluated by the remaining cell count present in the well measured via high content imaging (Figure 1F). These data indicate that accumulation of aberrant core protein induced apoptosis and that the compound itself in the absence of core did not induce this effect.

### CAM-A mediates anti-HBV activity in AAV-HBV transduced C57BL/6 mice

The anti-HBV activity of HAP-1 was tested *in vivo* in C57BL/6 mice transduced with 1x10^11^ viral genome equivalents AAV-HBV. Mice were orally gavaged once daily (QD) with 36 mg/kg HAP-1 or placebo for 28 days. Plasma sampling was performed frequently for the quantification of HBV DNA, HBsAg and ALT activity (Figure 2A). Treatment with HAP-1 induced a dose-related rapid decline of HBV DNA from a pre-treatment 4.2 log_10_ copies per µl to <LLOQ (2.08 log_10_ copies per µl) after 7 days of treatment; this suppression was maintained until the end of treatment on day 28. A continuous decline of HBsAg was observed from day 16 onwards in HAP-1 treated mice. The profound HAP-1-induced HBsAg decline was associated with a transient increase in ALT activity (Figure 2B, 2C), suggesting that treatment with CAM-A induced a biological effect in the liver, but not chronic toxicity, which is also supported by a lack of body weight loss (Figure 2D). Liver samples from three mice from the placebo and CAM-A treated group were analyzed at multiple time points to assess potential HAP-1 induced hepatic effects (Figure 2A). A time-dependent core and HBsAg reduction was observed in the liver of HAP-1 treated mice, indicating that treatment with HAP-1 induced the elimination of HBV-infected hepatocytes over time (Figure 2E). No effect on HBV DNA, HBsAg or ALT was observed in mice treated with placebo (Figure 2B, 2C).

**Figure 2.**
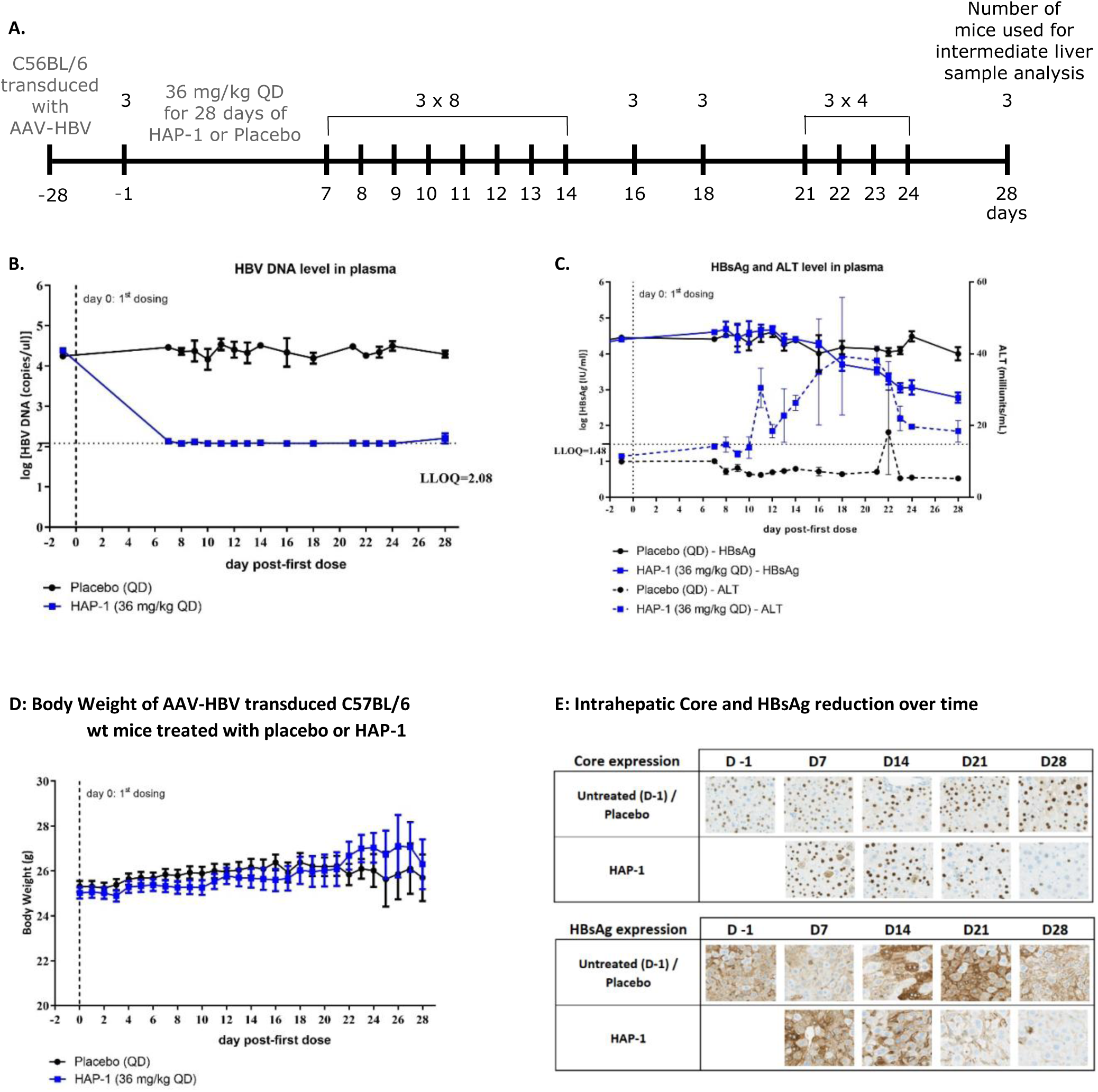
CAM-A anti-HBV activity in AAV-HBV transduced C57BL/6 wt mice. **(A)** C57BL/6 mice transduced with 1x10^11^ vg equivalents AAV-HBV were treated orally with placebo or HAP-1 (36 mg/kg QD) for 28 days. Time points and number of mice that were used for intermediate liver sample analysis are indicated. **(B)** Plasma sampling was performed frequently for the quantification of HBV DNA, HBsAg and ALT. **(C)** Body weight was monitored during treatment duration. **(D)** Immunohistochemistry for the detection of HBV core and HBsAg in liver of mice sacrificed at different time points are shown for placebo and HAP-1 treated mice for day -1, 7, 14, 21 and 28. QD once daily; LLOQ lower limit of quantification. In each graph data are shown as mean ± SEM and represent groups of 3 mice.

### Treatment with CAM-A induces apoptosis of infected hepatocytes in AAV-HBV-transduced mice

HAP-1 induces the formation and accumulation of intracellular aberrant core protein structures. To further understand how the induction of aberrant structures is associated with transient ALT increases and loss of core staining, we explored whether the accumulation of such aberrant subcellular structures may trigger cellular stress responses that result in the induction of apoptosis of HBV-infected hepatocytes *in vivo*. We applied a multiplex immunofluorescence analysis to liver slices from HAP-1- and placebo-treated mice at multiple time points for the detection of HBsAg, Ki67 (marker for proliferation) and DAPI (DNA stain). A TUNEL immunofluorescence stain for the detection of apoptotic cells that undergo extensive DNA degradation during the late stages of apoptosis was performed. An increase in apoptotic hepatocytes (up to ∼3.5%) was observed in HAP-1 treated mice compared to placebo treated mice (Figure 3A), which was accompanied by an increase in hepatocyte proliferation (determined by Ki67 measurement) (Figure 3B.1). At later time points (days 24 and 28), these proliferating hepatocytes were predominantly HBsAg negative (Figure 3B.2). The detection of apoptotic cells overlapped with the transient increase in ALT activity, suggesting that CAM-A-induced formation and accumulation of aberrant core protein in high level of core-expressing hepatocytes caused this effect. Consistent with the time-dependent HBsAg reduction in plasma we observed a time-dependent reduction of HBsAg positive hepatocytes in HAP-1 treated mice (Figure 3B.3).

**Figure 3.**
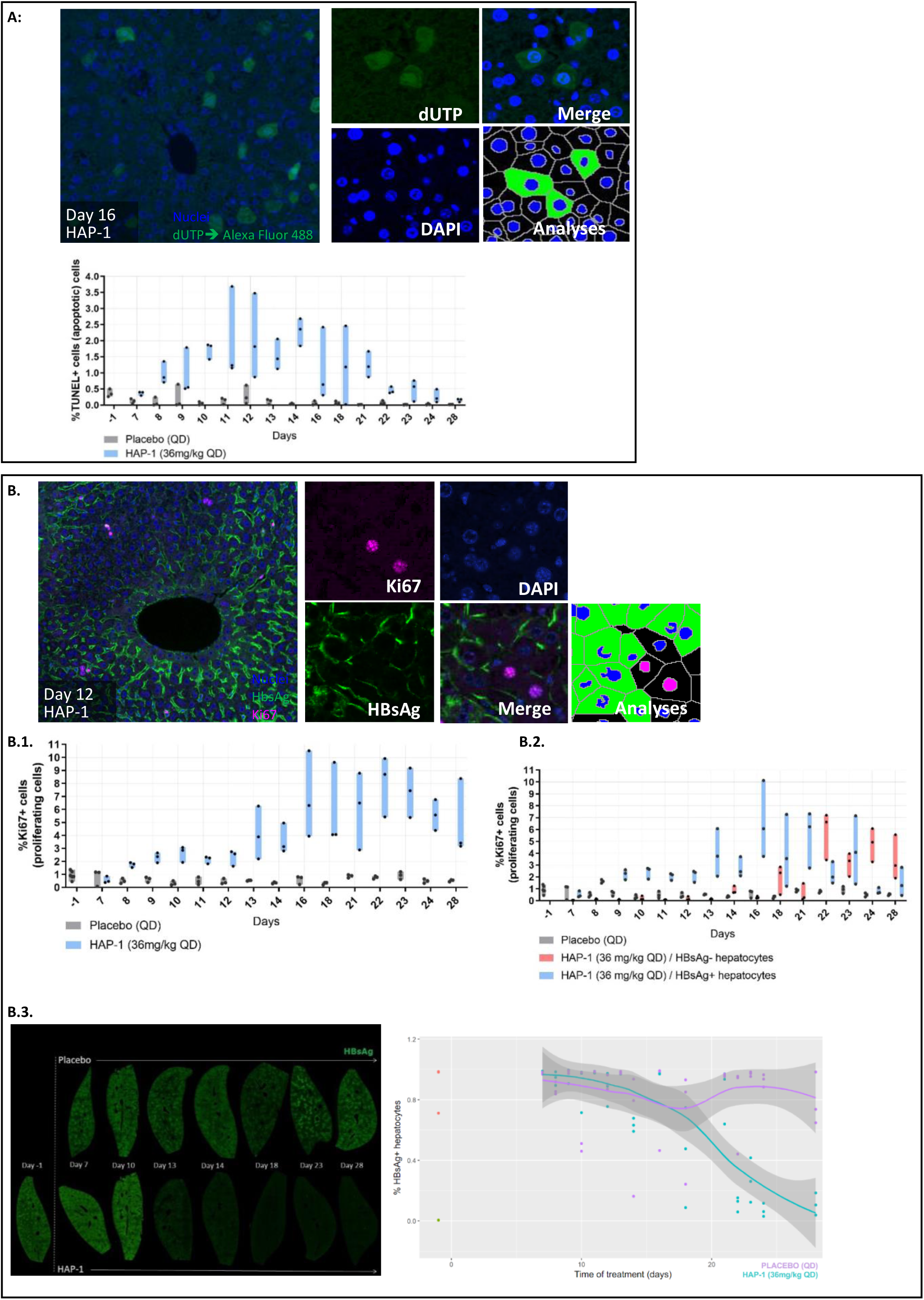
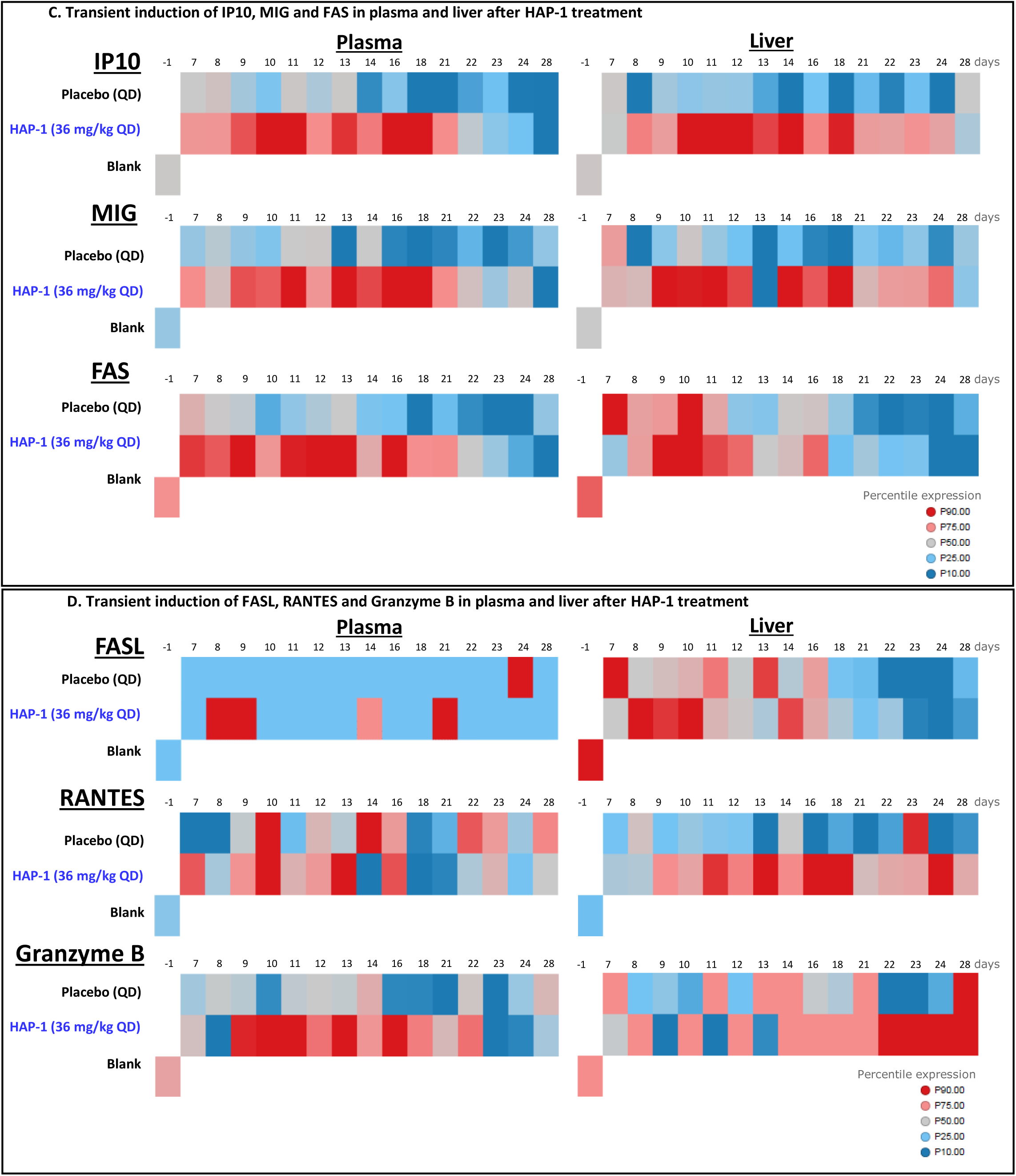
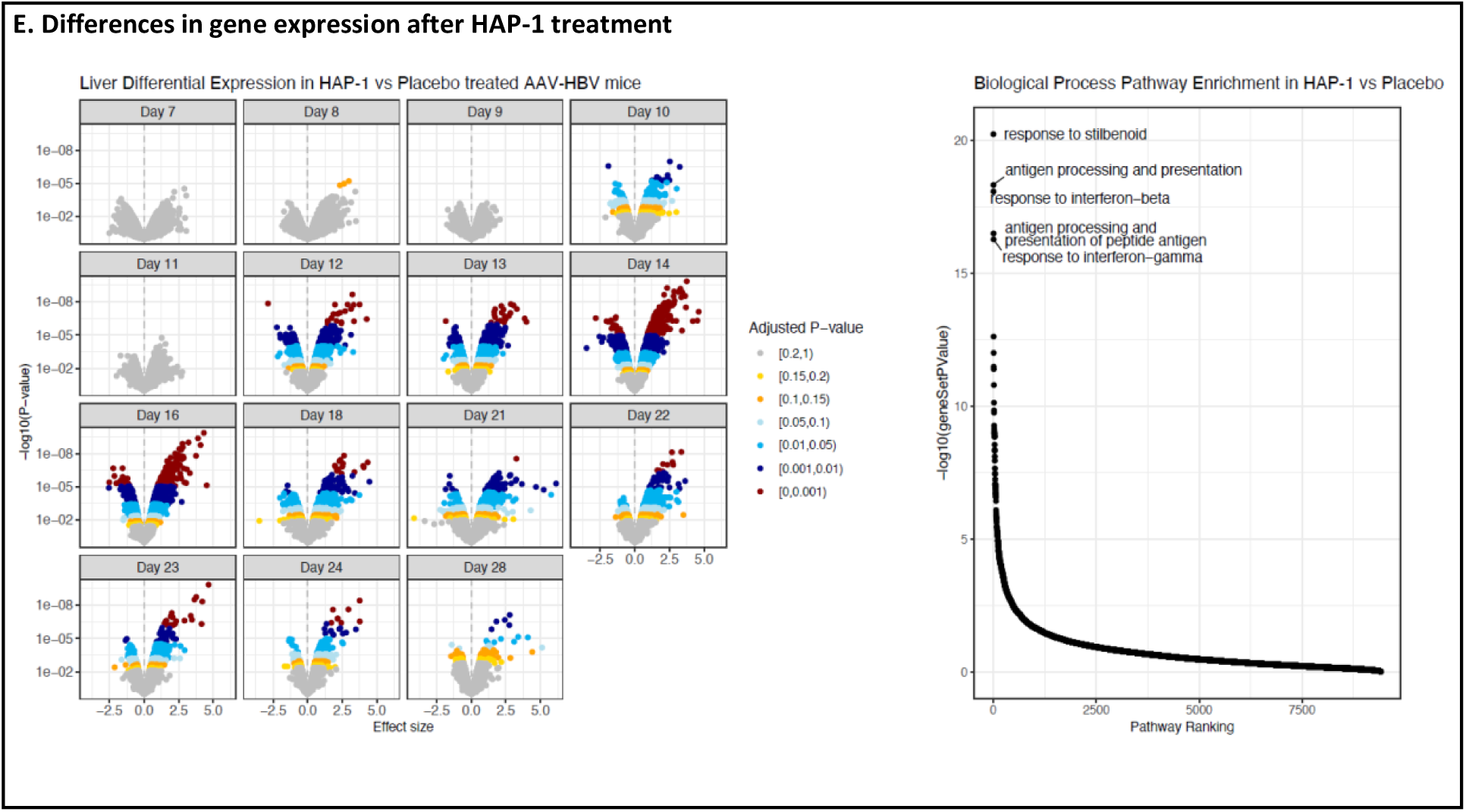
Treatment with CAM-A induces apoptosis and proliferation in the liver of AAV-HBV transduced mice. **(A)** Formalin-fixed, paraffin-embedded liver samples were sectioned at 5µm, mounted on SuperFrost Plus glass slides and used for TUNEL analysis. %TUNEL positive (apoptotic) cells are shown for HAP-1 (36 mg/kg QD) treated mice compared to placebo treated mice. **(B)** Alternative HBsAg/Ki67/Cleaved caspase 3/DAPI stained images are shown with the calculation of %Ki67+ cells representing proliferating cells. A subdivision between HBsAg- and HBsAg+ cells of the Ki67+ cells for HAP-1 treated AAV-HBV mice is shown as well. Time dependent immunofluorescence staining against HBsAg (green) on liver slices with the correlating calculation of % HBsAg+ hepatocytes over time of HAP-1 treated AAV-HBV mice versus placebo treated. For A and B, data are shown as floating bars indicating the range and the mean by black dots. **(C-D)** Heatmaps from mouse plasma samples and liver homogenates for IP10, MIG, FAS, FASL, RANTES and Granzyme B. Percentile average expression is shown. **(E)** Volcano plots per each time point for log-fold change difference expression of genes between placebo and HAP-1 treated AAV-HBV mice, coloured by adjusted fdr p-value. Pathway ranking plot from gene set enrichment analysis between placebo and HAP-1 treated AAV-HBV mice. Top 5 pathways, with lowest p-value score are shown. QD once daily; % percentage. The data adds on to the in vivo study explained in figure 2.

### Treatment with CAM-A transiently induces immune activation

To assess if additional factors contribute to the CAM-A mediated clearance of HBV-infected hepatocytes, mouse plasma samples and liver homogenates were analyzed for the presence of immune mediators using the Mouse Luminex® Discovery Assay. A transient increase of IP10, MIG and FAS (increase was less clear for FAS) was observed in plasma and liver homogenates of HAP-1 treated mice (Figure 3C), but not in mice treated with vehicle only. FASL was only detected in plasma, whereas RANTES and Granzyme B were only detected in liver homogenates (Figure 3D). No evident HBV antigen specific T cells were detected in spleen by ex vivo IFN-γ ELIspot on day 7, 16, 21 and 28 post dosing (data not shown).

### Effect of CAM-A treatment on liver gene expression

To understand whether changes in RNA expression were observed during the treatment-induced clearance of HBV-infected hepatocytes, liver gene expression was analyzed at different time points. Differentially expressed genes between HAP-1 treated versus vehicle-treated mice were mainly observed between day 12-16, with over 500 genes significantly expressed (Figure 3E, Supplementary data table 1). Days 14 and 16 showed 1,947 and 1,743 significantly expressed genes, respectively (adjusted p-value threshold of 0.05), indicating profound transcriptional changes. Pathway analysis showed an enrichment in processes involved in antigen processing and presentation and responses to interferon. Increased gene expression for *Cd74*, *H2-Ab1*, *H2-Eb1*, *H2-Q7* was observed from day 13 to 16, coinciding with the appearance of apoptotic events.

### CAM-A mediates anti-HBV activity in AAV-HBV transduced C57BL/6 SCID mice lacking T and B cells

The cytokine and microarray data from studies in C57BL/6 mice suggested that adaptive immune responses might contribute to the elimination of HBV infected hepatocytes and the accompanying reduction in HBsAg. To study this in more detail, C57BL/6 SCID mice that lack mature B and T lymphocytes, were transduced with 1x10^11^ viral genome equivalents AAV-HBV and treated orally for 28 days once daily (QD) with 36 mg/kg HAP-1 or placebo (Figure 4A). There was no apparent difference in HBV DNA (Figure 4B), HBeAg (Figure 4C) and HBsAg (Figure 4D) levels at the beginning of dosing between C57BL/6 and C57BL/6 SCID AAV-HBV transduced mice showing that the efficiency of transduction was similar in both mouse strains. Plasma HBV DNA levels decreased to reach the LLOQ after 7 days of dosing in C57BL/6 and C57BL/6 SCID (Figure 4B), indicating that the compound was equally efficacious in suppressing HBV replication in both mouse strains. However, there was an apparent difference observed in time and extent between C57BL/6 and C57BL/6 SCID transduced mice for both HBsAg decline and transient increase in ALT (Figure 4D). A delayed and less pronounced decline of HBsAg and a delayed transient increase in ALT was observed in C57BL/6 SCID compared to C57BL/6 mice (Figure 4D). Also, a delayed and less pronounced HBeAg decline was observed in C57BL/6 SCID mice (Figure 4C).

**Figure 4.**
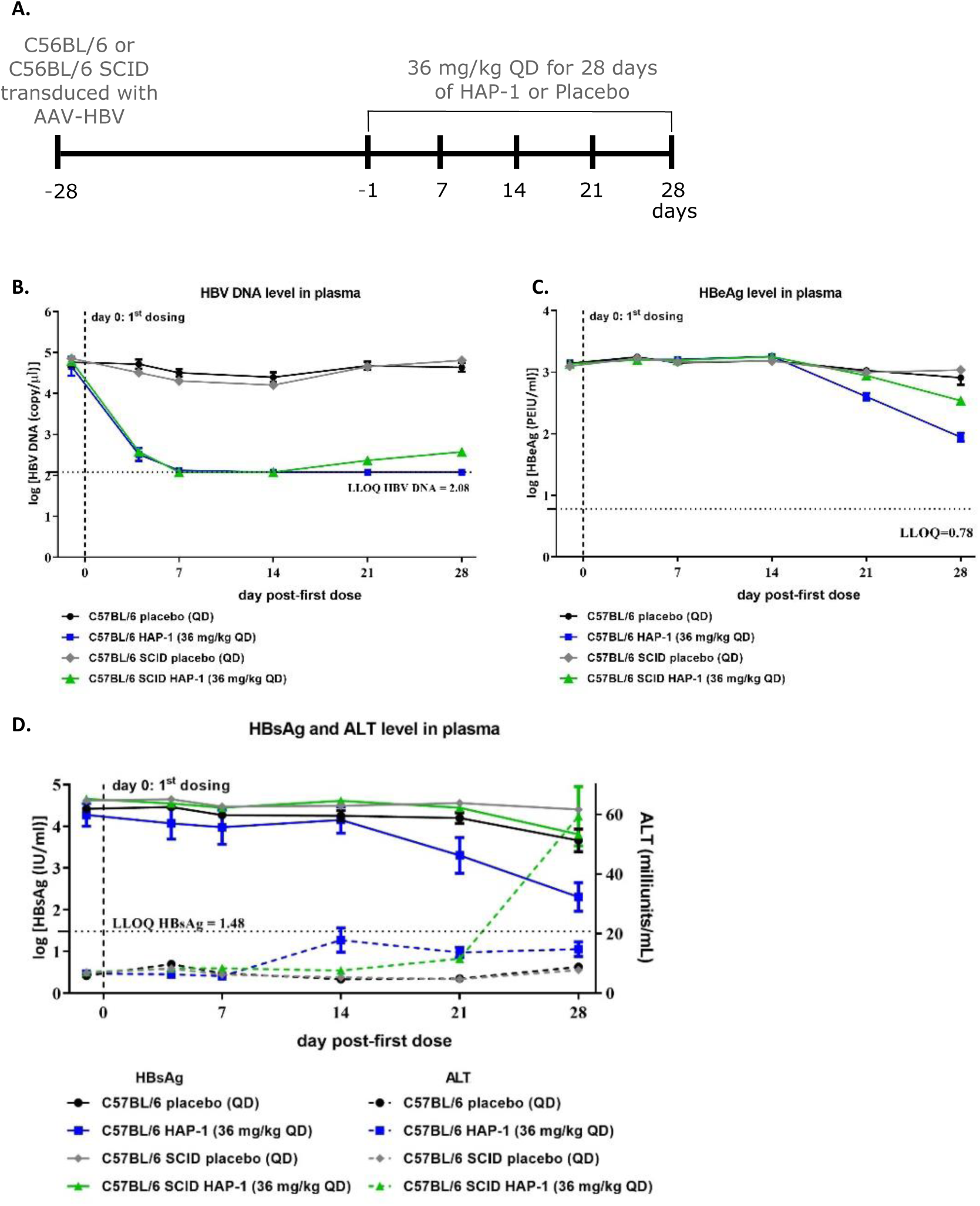
HBV DNA, HBeAg, HBsAg decline and transient increase in ALT in C57BL/6 and C57BL/6 SCID mice after HAP-1 treatment. **(A)** C57BL/6 and C57BL/6 SCID mice that lack mature B and T lymphocytes, were transduced with 1x10^11^ vg equivalents AAV-HBV and treated orally for 28 days with HAP-1 (36 mg/kg QD) or placebo. **(B-C)** Plots represent HBV DNA and HBeAg levels in plasma during treatment duration. **(D)** Correlation plot is shown between HBsAg levels and ALT increase in plasma during treatment duration. QD once daily; vg viral genomes; LLOQ lower limit of quantification. Data are shown as mean ± SEM and represent groups of 5 mice.

### CD8 T cells are dispensable for CAM-A mediated anti-HBV activity in AAV-HBV transduced C57BL/6 mice

Results from a study in AAV-HBV transduced C57BL/6 SCID mice suggested that adaptive immune responses contribute to clearance of HBV-infected hepatocytes. To study if CD8+ T-cells contribute to the observed effect, C57BL/6 AAV-HBV transduced mice had their CD8+ T-cells depleted with an anti-CD8 antibody, or were treated with an isotype control antibody, and additionally dosed orally (gavage) once daily (QD) with placebo or 20 mg/kg HAP-2 for 56 days (Figure 5A). Successful depletion of CD8+ T-cells, and maintained over time, was confirmed by FACS (Figure 5B). Treatment with 20 mg/kg HAP-2 lead to a similar response for HBsAg and HBeAg, in both CD8+ depleted and non-depleted mice suggesting that CD8+ T-cells are dispensable for the clearance of HBV-infected hepatocytes (Figure 5C). However, it cannot be excluded that other components of the adaptive immune system are involved.

**Figure 5.**
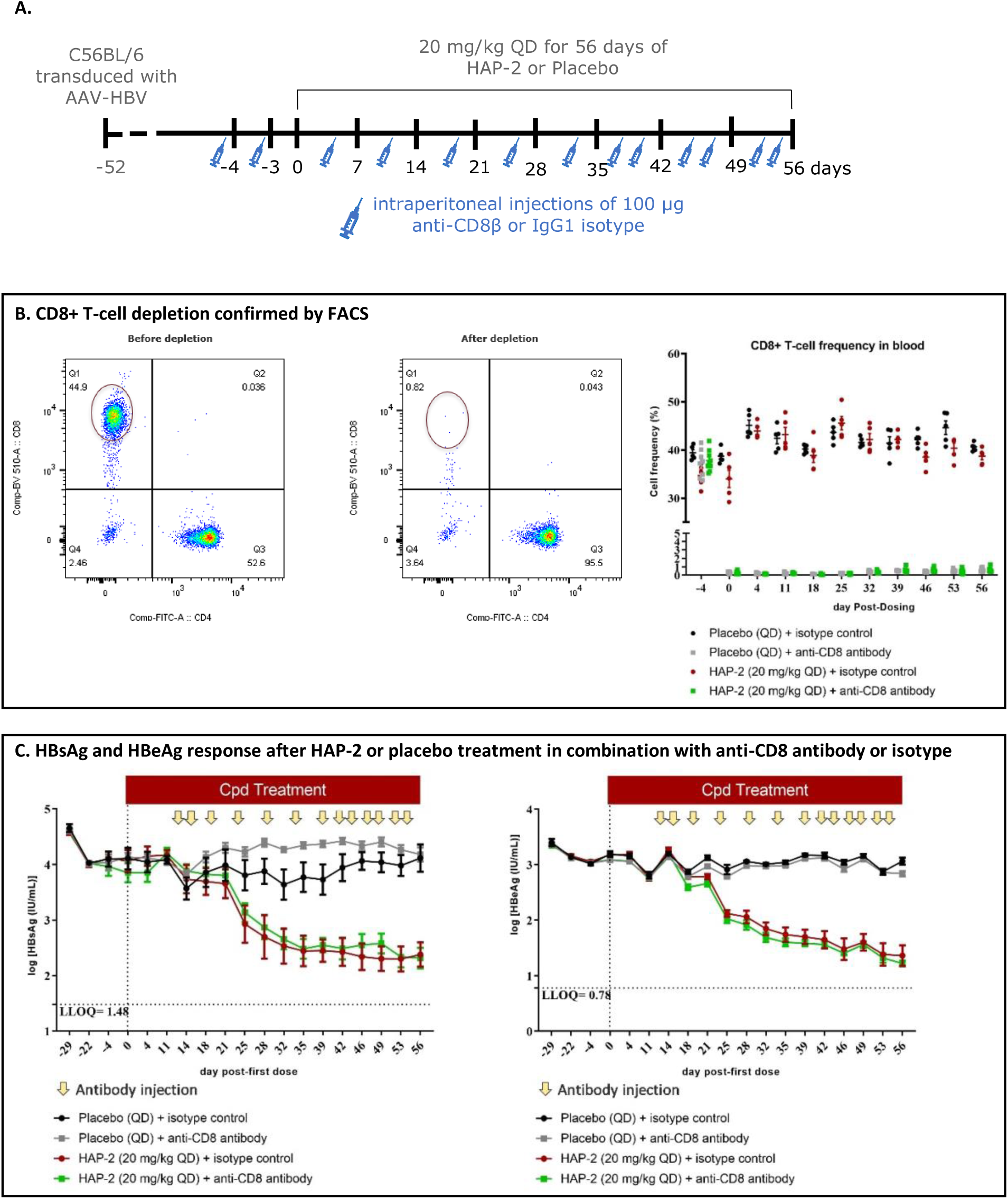
CAM-A anti-HBV activity in AAV-HBV transduced C57BL/6 wt mice with depleted CD8+ T-cells. **(A)** AAV-HBV transduced C57BL/6 mice with 1x10^11^ vg equivalents were treated orally with placebo or HAP-2 (20 mg/kg QD) for 56 days. CD8+ T-cell depletion, intermediate plasma sampling, sacrifices and number of mice that were sacrificed are indicated on the timeline. **(B)** FACS analysis gating for CD8+ T-cells with representation of CD8+ T-cell frequency in blood of HAP-2 treated HBV-AAV mice versus placebo treated. **(C)** HBsAg and HBeAg response after treatment with HAP-2 versus placebo in both CD8+ depleted and non-depleted mice. QD once daily; LLOQ lower limit of quantification. Data are shown as mean ± SEM and represent groups of 5 mice for the isotype treated control groups or 8/9 mice for the anti-CD8 treated groups.

### Induced elimination of HBV-infected hepatocytes is dependent on high viremia and antigen expression

The AAV-HBV mouse model represents a single round infection model, as HBV cannot infect mouse hepatocytes (27, 28) nor can mouse hepatocytes support the formation of authentic cccDNA (29) and the transcription of HBV RNAs is dependent on the AAV-HBV template within the transduced hepatocytes. Therefore, the transduction of mice with different AAV-HBV doses results in different levels of viremia and antigen expression in the hepatocytes. To study this core protein dependency in more detail, we transduced mice with four different doses of AAV-HBV (5x10^9^, 1x10^10^, 2.5x10^10^ and 1x10^11^ AAV-HBV vg equivalents) and treated orally for 56 days once daily (QD) with 36 mg/kg HAP-1 or placebo (Figure 6A). An inoculum-dependent increase in HBV DNA and HBeAg (supplementary figure 1) as well as HBsAg (Figure 6B) and HBcrAg (Figure 6C) levels was observed in mice transduced with 5x10^9^, 1x10^10^, 2.5x10^10^ and 1x10^11^ vg equivalents respectively, indicating that viremia is dependent on the amount of AAV-HBV inoculum used for transduction and not on spread of infection. Treatment with HAP-1 reduced HBV DNA levels to LLOQ (supplementary figure 1). HBsAg was reduced over time only in mice transduced with 2.5x10^10^ and 1x10^11^ AAV-HBV vg equivalents (Figure 6B). An earlier and steeper decline of HBsAg was observed in mice transduced with 1x10^11^, suggesting that higher levels of viremia / expression of core antigen in hepatocytes led to a faster and greater accumulation of aberrant core protein resulting in an earlier and stronger induction of apoptosis upon treatment. In addition, higher levels of HBcrAg and a steeper decline upon treatment were observed in mice transduced with 1x10^11^ vg equivalents (Figure 6C). No significant effect on HBsAg and HBcrAg was observed in mice transduced with 5x10^9^ or 1x10^10^ vg equivalents. Consistent with the HBsAg observations, a more pronounced and earlier transient increase in ALT activity was observed in mice transduced with 1x10^11^ vg equivalents compared to mice transduced with 2.5x10^10^ vg equivalents upon treatment (Figure 6D). No transient increase in ALT activity was observed in mice transduced with 5x10^9^ and 1x10^10^ vg equivalents nor in any of the placebo-treated mice (Figure 6D).

**Figure 6.**
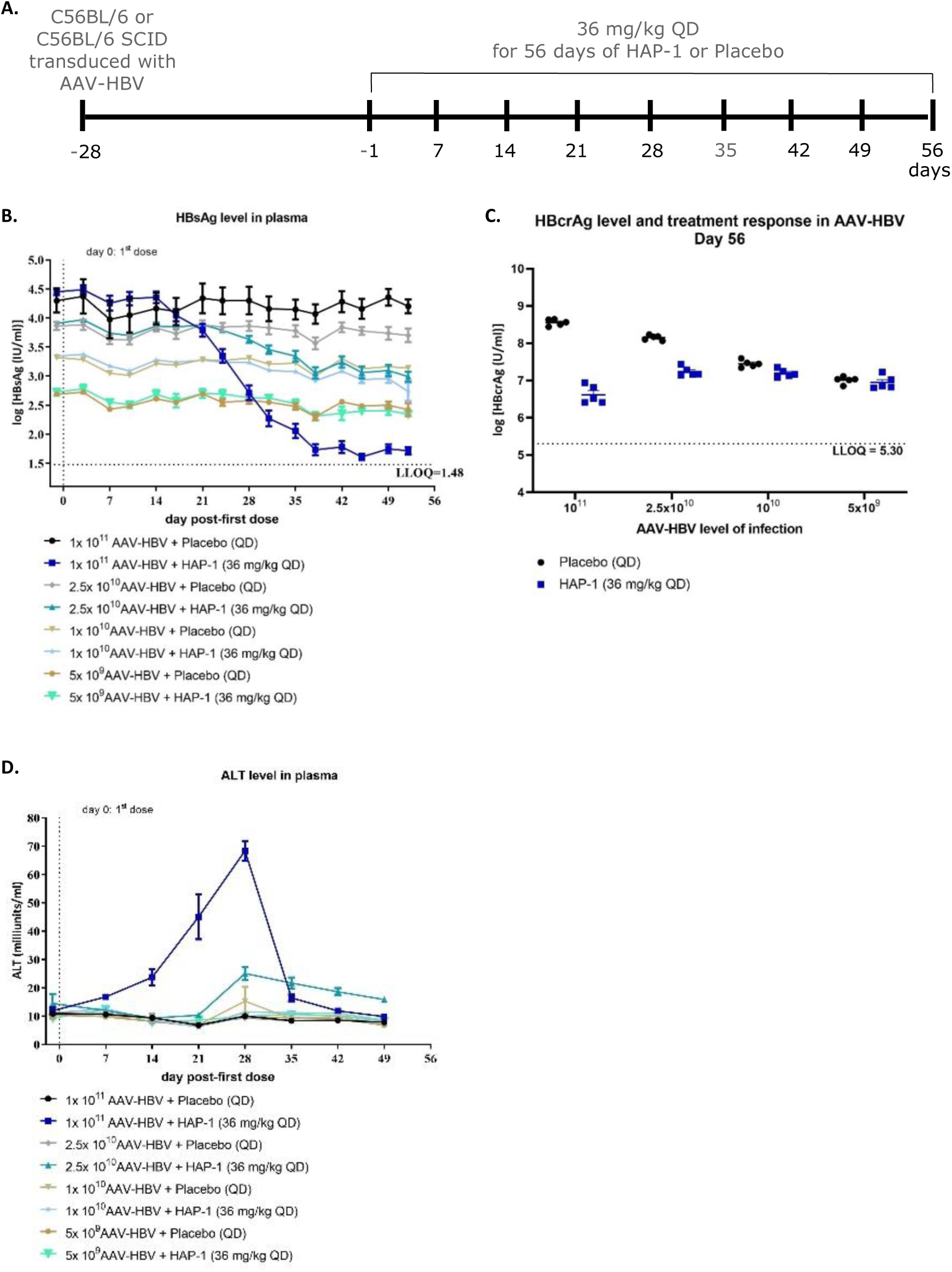
HBsAg and HBcrAg levels in mice transduced with different amounts of AAV-HBV. **(A)** C57BL/6 mice transduced with four different amounts of AAV-HBV (5x10^9^, 1x10^10^, 2.5x10^10^ and 1x 10^11^ AAV-HBV vg equivalents) were treated orally with placebo or HAP-1 (36 mg/kg QD) for 28 days. **(B-D)** Plots represent HBsAg, HBcrAg and ALT levels in plasma during treatment duration. QD once daily; vg viral genomes; LLOQ lower limit of quantification. Data are shown as mean ± SEM and represent groups of 5 mice for the placebo treated control groups and 10 mice for the HAP-1 treated groups.

### CAM-A induced elimination of HBV-infected hepatocytes is dependent on *de novo* core protein translation

Given that the CAM-A induced clearance of HBV-infected hepatocytes is dependent on high viremia and high core protein levels, we next investigated if the induction of apoptosis was dependent on *de novo* core protein translation. To do this a GalNac-conjugated HBV siRNA was used in combination with the CAM-A inhibitor in the AAV-HBV mouse model. C57BL/6 mice were transduced with 1x10^11^ vg equivalents AAV-HBV and treated with placebo, HAP-3 (12 mg/kg oral for 12 weeks daily QD), siRNA (3 mg/kg s.c. four times every three weeks) and a combination of HAP-3 with siRNA, followed by a 12-week washout (Figure 7A). Plasma samples were collected at multiple time points throughout the study to assess the effect on HBV DNA, HBeAg, HBsAg, HBcrAg and ALT (Figure 7B-F). HBV DNA levels declined to, and remained at, LLOQ during HAP-3 monotherapy and during washout, indicating that HAP-3 monotherapy eliminated HBV-infected hepatocytes as observed previously (Figure 7B).

**Figure 7.**
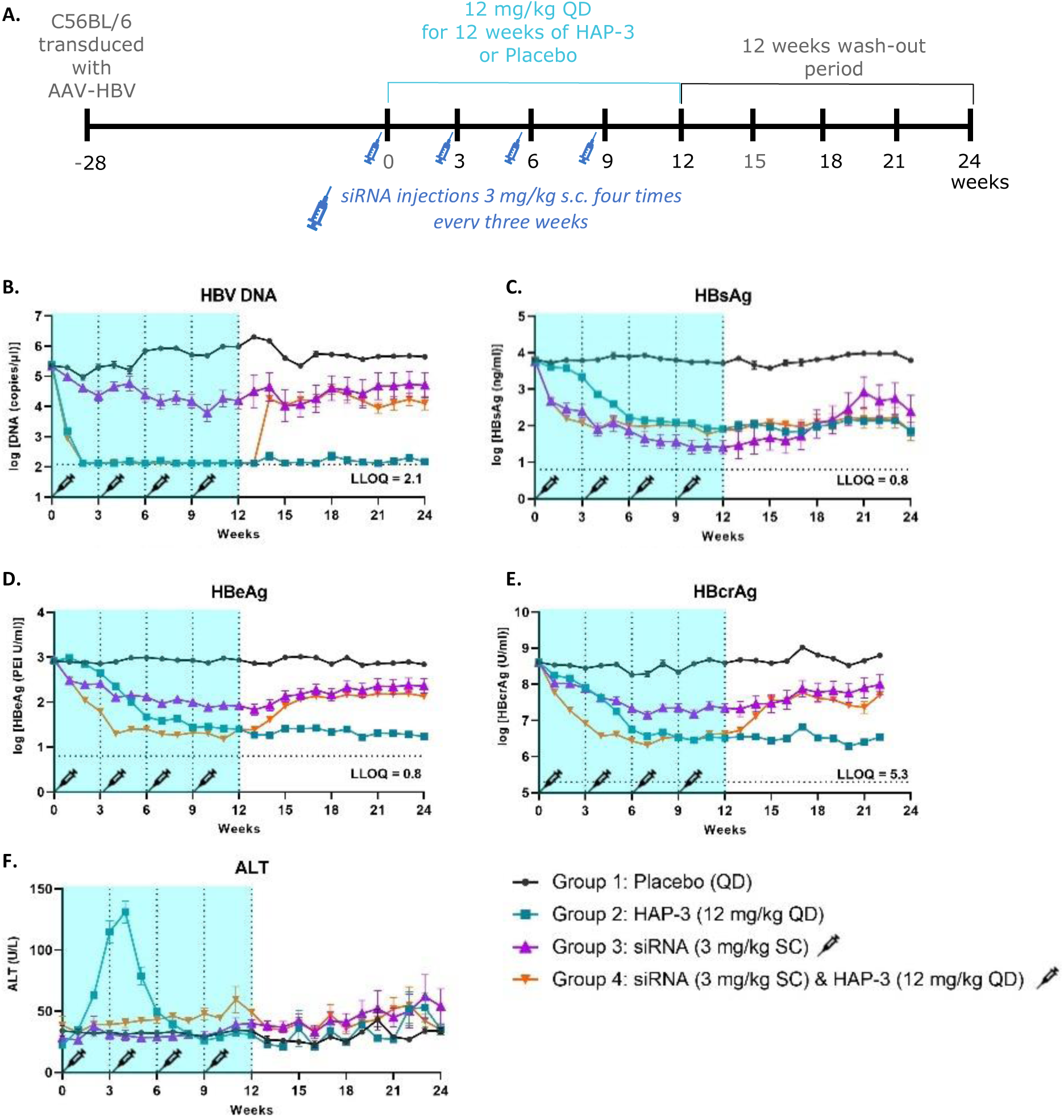
CAM-A induced elimination of HBV-infected hepatocytes is dependent on de novo core protein translation. **(A)** C57BL/6 mice were transduced with 1x10^11^ vg equivalents AAV-HBV and treated with placebo, HAP-3 (12 mg/kg oral for 12 weeks daily QD), siRNA (3 mg/kg s.c. four times every three weeks) and a combination of HAP-3 with siRNA, followed by a 12-week washout. **(B-F)** Plasma samples were collected at multiple time points throughout the study to assess the effect on HBV DNA, HbeAg, HbsAg, HbcrAg and ALT. QD once daily; vg viral genomes; s.c. subcutaneous; LLOQ lower limit of quantification. Data are shown as mean ± SEM and represent groups of 5 mice.

Treatment with siRNA alone induced a less pronounced reduction of HBV DNA than HAP-3 and a plateau around week six prior to the third siRNA dose. Additional siRNA administration did not reduce the HBV DNA further. Mean HBV DNA levels remained constant during the washout phase, likely due to a long half-life of the siRNA. However, a larger level of variation across mice was observed (increased standard errors on the derived means during the washout compared to the treatment phase). The combination of siRNA with HAP-3 induced a rapid decline of HBV DNA to the LLOQ. A rebound of HBV DNA to levels observed with siRNA monotherapy occurred after 2 weeks washout (week 14), indicating that antiviral activity was mediated by the siRNA and not by HAP-3. Rebounds were also observed for HBeAg and HBcrAg in the HAP-3 + siRNA treatment group during the wash out phase whereas no rebound was observed in the HAP-3 monotherapy group, suggesting that combination treatment with siRNA interfered with the HAP-3-induced clearance of HBV-infected hepatocytes (Figure 7D,7E). This assumption is supported by the observation that a transient increase in ALT activity was only observed in the HAP-3 monotherapy group and not in the HAP-3 plus siRNA combination treatment group (Figure 7F). Treatment with siRNA alone or in combination with HAP-3 resulted in a faster and steeper HBsAg decline than HAP-3 treatment alone. A rebound was observed during the washout phase (Figure 7C). Altogether these data suggest that CAM-A induced clearance of HBV-infected hepatocytes is dependent on *de novo* core protein translation.

### CAM-A modulation mediate anti-HBV activity in HBV-infected chimeric mice with humanized liver

To study the anti-HBV activity of CAM-A in a model with “authentic” HBV infection, HBV infected chimeric humanized liver mice were treated orally for 84 days once daily (QD) with 36 mg/kg HAP-1 or placebo. Approximately four times lower compound concentrations were observed in the liver of chimeric mice compared to C57BL/6 mice treated with the same dose (data not shown).

Repeated serum sampling was performed to quantify HBV DNA, HBe-, HBsAg, HBcrAg, hAlb and hALT activity. Mice samples were taken at multiple time points during treatment (see Figure 8A) to assess HAP-1 induced effects in the liver. Treatment with HAP-1 induced a rapid decline of HBV DNA, indicating that the compound suppressed HBV replication (Figure 8B). In contrast to what was observed in AAV-HBV transduced mice, a clear HBsAg and HBcrAg reduction was only observed from day 35 onwards and the decline was less steep. Interestingly, HBsAg decline was also associated with a transient increase in human ALT activity (Figure 8B), suggesting that treatment with CAM-A induced a transient biological effect in the human hepatocytes. Stable human albumin levels were observed over the entire treatment duration (Figure 8C) and no body weight loss (Supplementary figure 2) was observed during HAP-1 treatment, further indicating that the compound itself is unlikely to cause cytotoxic effects.

**Figure 8.**
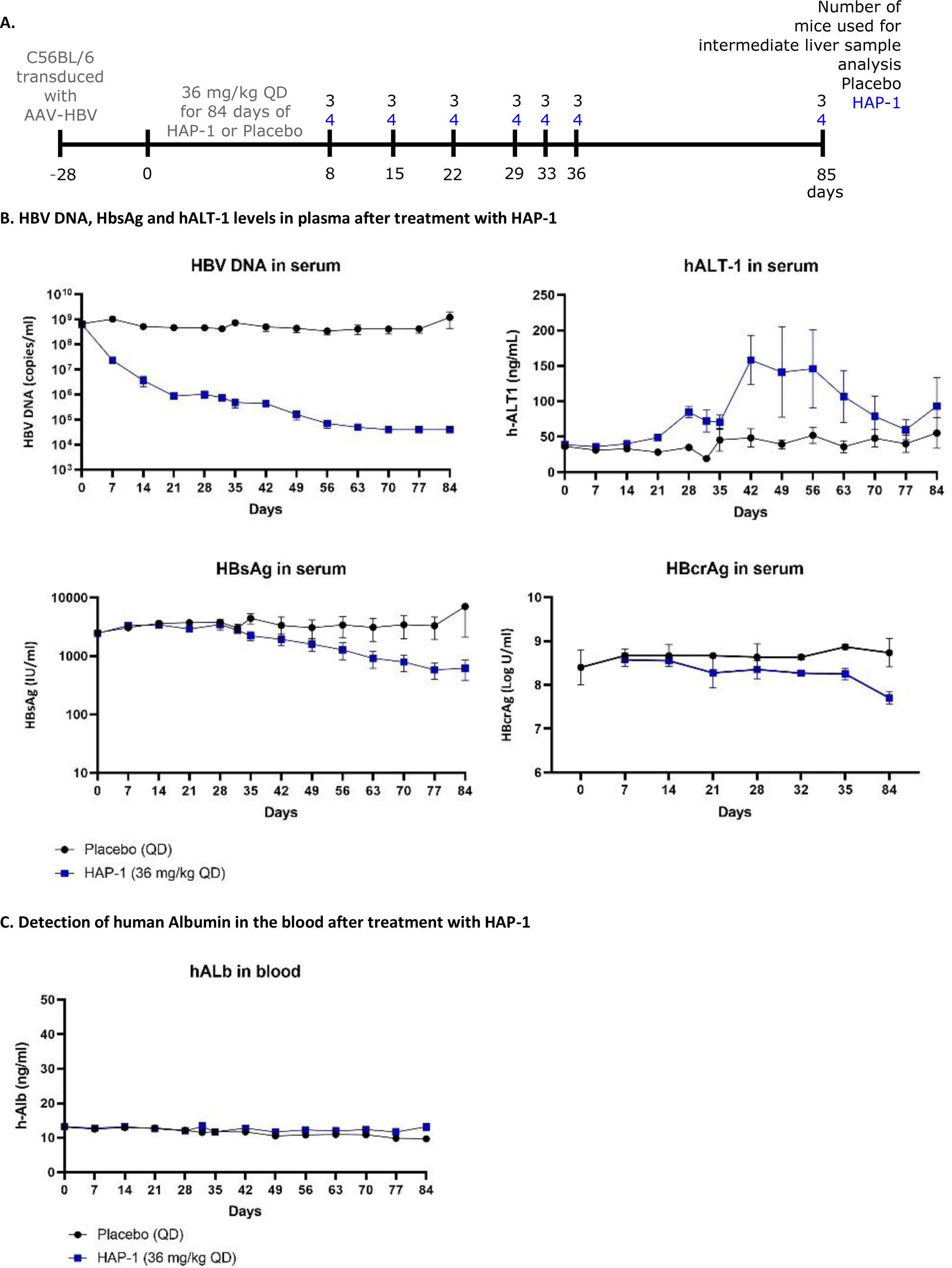
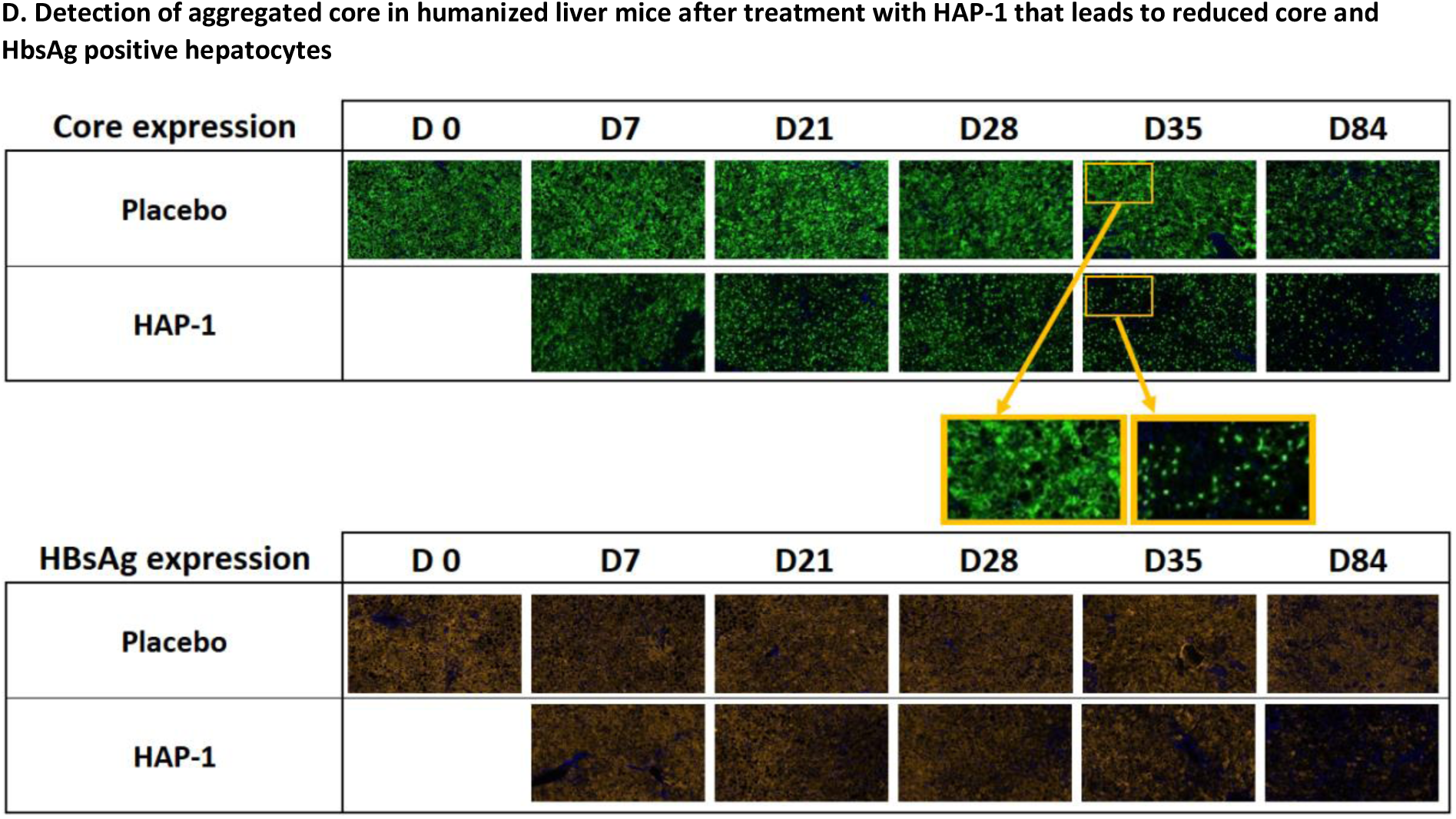
CAM-A anti-HBV activity in HBV-infected chimeric humanized liver mice. **(A)** HBV infected chimeric humanized liver mice were treated orally for 84 days once daily (QD) with 36 mg/kg HAP-1 or placebo. Time points and number of mice that were used for intermediate liver sample analysis are indicated. **(B)** Serum sampling was performed frequently for the quantification of HBV DNA, HBsAg, HBcrAg and hALT-1 activity. **(C)** Detection of human albumin levels in blood during treatment duration. **(D)** Core and HBsAg detection over time in the liver of mice treated with HAP-1 or placebo with a more detailed image of the core staining on day 35 to visualize the formation of aberrant core structures after HAP-1 treatment versus the normal core detected in the placebo treated condition. QD once daily. Data are shown as mean ± SEM and represent groups of 3 mice for the placebo treated control groups and 4 mice for the HAP-1 treated groups.

Aberrant core protein, represented by “dot-like structures”, was detected in the liver of mice treated with HAP-1 and a reduction of core and HBsAg positive hepatocytes was observed over time (Figure 8D). We were not able to detect apoptosis and proliferation of hepatocytes in the liver of HAP-1 treated mice in contrast to the observations in the AAV-HBV mouse model.

## DISCUSSION

HBV represents a global health burden that necessitates the development of new regimens for CHB, aimed at achieving increased functional cure rates after finite duration therapy. At a first glance, targets for direct-acting antivirals may be considered exhaustive as the condensed HBV genome only encodes seven proteins (30). Yet, all these viral proteins fulfill multiple roles in the HBV life cycle, which allows for broader pharmacologic considerations when interfering with their functions. HBV core protein provides an attractive target for the development of new interventions because it acts at multiple key steps during viral replication (11). In this study, we evaluated the anti-HBV activity and mode of action *in vitro* and *in vivo* of heteroaryldihydropyrimidines (HAPs), capsid assembly modulators classified as CAM-A. HAPs are known to induce heterogeneous forms of capsid in vitro (14). This study is consistent with previous reports showing that treatment with HAPs CAM-A modulators induce the aggregation of HBV core protein in an inducible HBV replicating cell line (26). Accumulation of such aberrant protein structures was shown to lead to apoptosis of the cells *in vitro* when cultured for an extended period in presence of compound. This was not an artifact since a dose and time dependent increase in caspase 3/7 activity was completely absent in cells treated with compound but lacking core expression. The observation that apoptosis is not immediately evident led us to hypothesize that the effects induced by these aggregated core protein structures over time may be the result of an accumulation of the unfolded protein response or UPR. The UPR functions to restore the normal homeostasis of a cell when stressed by the accumulation of misfolded proteins. If this response cannot adequately cope with the insult within a certain time span, the cell undergoes apoptosis (31). Similar observations of aggregate induced effects by HAPs have been described by Huber *et al*. (32). The authors report on the association of core aggregates with promyelocytic leukemia nuclear bodies or PML-NBs in infected cells when exposed to CAM-A modulator HAP Bay38-7690. Although the publication does not describe the induction of apoptosis, Huber et al do suggest that this accumulation of core protein with PML-NBs could potentially disrupt cellular process and impart cell death. A recent study from Kum *et al.* showed in the AAV-HBV model -A induced aggregation and apoptosis (33).

Notably, when transduced mice were dosed with HAPs, not only did they profoundly suppress HBV replication but the treatment also led to the reduction of HBsAg accompanied by a transient increase in ALT in absence of any safety signals. This decline in HBsAg has not been described for compounds belonging to the CAM-E class pointing towards an additional differentiating property of CAM-As. Furthermore, this ALT increase was absent in naïve mice dosed with compound. In examinations of the liver of AAV-HBV transduced mice receiving compound, it was apparent that compared to placebo treated animals mice treated with HAPs showed a time-dependent decline in core and HBsAg expressing hepatocytes coinciding with the increase in ALT. To connect the apoptosis observed *in vitro* with a potential similar event *in vivo*, we performed a multiplex immunofluorescence on the liver looking at HBsAg, DNA, and Ki67, a known marker for proliferation. These results showed that CAM-A treatment increased the percentage of apoptotic cells along with an enhanced proliferation of HBsAg negative cells with a subsequent reduction in HBsAg positive cells. The accumulation of aberrant core structures within hepatocytes by HAPs resulted in cell death and was represented by a transient ALT increase and repopulation of the liver by non-infected hepatocytes.

In exploiting the single-round infection characteristics of the AAV-HBV mouse model, we also demonstrated that the size of the inoculum determines the level of antigen expressed in the transduced hepatocytes and in turn the extent of decline in viral antigen and ALT activity upon treatment with CAM-A. No CAM-A mediated antigen reduction and transient increases in ALT were observed in mice with low(er) levels of viremia, implying that a steady state between CAM-A induced formation of aberrant core protein structures and their degradation in hepatocytes of those mice could be maintained. In contrast, CAM-A treatment induced a faster and steeper decline of antigen in mice with the highest level of viremia suggesting that faster accumulation of aberrant core protein led to an earlier induction of apoptosis. Our titration studies in transduced mice suggest that HBcrAg levels above 8 log10 IU/ml would be required to observe a CAM-A mediated HBsAg decline. High HBcrAg levels above 8 log10 U/ml are mainly observed in naïve HBeAg+ patients and HBcrAg exhibits good correlation with intrahepatic cccDNA (34). Therefore, it is likely that HBsAg reductions during CAM-A treatment would only be observed in treatment naïve, HBeAg+ and especially immunotolerant patients who have the highest HBcrAg levels in the periphery and high core protein levels in hepatocytes.

This is corroborated by the CAM-A / siRNA combination study data that showed the CAM-A mediated effect of HBV antigen reduction is dependent on the de novo synthesis of core protein and that preexisting core protein levels are insufficient to induce apoptosis.

Given that transduced mice do not recapitulate the full replication cycle of HBV and to exclude that this model therefore harbors a bias towards this pharmacodynamic effect of a CAM-A HAP modulator, we also conducted a study in chimeric mice with a humanized liver. These mice can be infected with HBV and the level of viremia is dependent on de novo infection and viral spread. Consistent with the observations in vitro, the formation of aberrant core protein structures into “dot-like structures” was also apparent in hepatocytes of chimeric mice treated with CAM-A, demonstrating that similar pharmacological effects occur in human hepatocytes *in vivo*. This model also exhibited an HBsAg decline and transient increase in human ALT albeit delayed and less pronounced compared to transduced mice suggesting that a similar biological effect was induced. However, we were not able to observe apoptosis and proliferation, likely due to too early and infrequent sampling in the study. The difference between the two mouse models might be attributed to an approximately lower compound exposure in chimeric mice compared to AAV-HBV mice at the same dose. In addition, chimeric mice do not possess an adaptive immune system and we cannot exclude that the delays and reduced extent of HBsAg decline, and ALT are due to an immunocompromised background.

To elaborate further on the potential role of adaptive immunity in response to CAM-A treatment, we compared the antiviral effects in both transduced wild type and SCID mice and evaluated the response of HAP treatment in transduced mice depleted in CD8+ T-cells. These studies collectively showed that CD8+ T cells are not required for CAM-A mediated HBsAg declines. No HBV specific T cells were detected in the spleen suggesting that CAM-A mediated induction of HBV-infected hepatocyte cell death is unlikely to generate potent immunogenic responses to control the virus and eliminate infected cells. However, it cannot be excluded that other factors beyond the induction of apoptosis contribute to the observed effects as we observed differentially expressed genes in the AAV-HBV mouse model before the ALT flare leading up to the HBsAg decline. Gene set enrichment showed that the upregulated genes were involved in antigen presentation and major histocompatibility complex class II. Interestingly, CD74 was upregulated and is known as a receptor for MIF mediated signaling in tissue repair (32). In liver, this same MIF-CD74 signaling has shown a protective role in nonalcoholic fatty liver disease (33). All these observations at the RNA level suggest that some component of the adaptive immune responses might play a role in the clearance of HBV-infected hepatocytes although it is not clear what it is and to what extent it is directly contributing or collateral to the induced cell death. This is also supported by the transient induction of cytokines that can be linked to adaptive immune responses (e.g. RANTES and Granzyme B). However, it also cannot be excluded that the detection of certain cytokines is a response towards the induction of apoptosis and the clearance of apoptotic cells.

The immunostimulatory effect of a CAM-A (BAY41-4109) had been evaluated in vitro (35). Direct treatment of HBV-infected PHH did not stimulate the production of type I and type III IFN. However, treatment enhanced the antiviral ISG expression induced by IFN-α in HBV-infected PHHs. Our AAV-HBV mouse data showed a transient induction of inflammatory response in the liver which was associated with a transient ALT increase, HBsAg decline and elimination of infected hepatocytes. However, there was no induction of type I IFN in liver by CAM-A treatment. These data suggests that treatment with CAM-A alone is not sufficient to stimulate potent immune response to control HBV infection in AAV-HBV transduced mice. In summary, we demonstrated that CAM-A treatment eradicates some HBV infected hepatocytes in preclinical animal models through selective induction of apoptotic processes, and that this effect is dependent on de novo translation and high levels of core antigen. HBV core protein synthesis is dependent on the presence of cccDNA (13) and as a result, CAM-A treatment would only lead to the specific eradication of HBV-infected hepatocytes with transcriptionally active cccDNA, leaving hepatocytes with silenced cccDNA and/or integrated HBV DNA untouched. Meaningful HBsAg reductions have not been observed in clinical trials with CAM-A so far (24). However, increases in ALT activity in subjects with high HBcrAg levels treated with RO7049389 were reported (22). Since subjects were only treated for up to 4 weeks it cannot be excluded that HBsAg reductions would have occurred during longer treatment as observed in our chimeric mouse study.

## Abbreviations

A full list of abbreviations can be found as supplementary data file.

## Supporting information

Abbreviations

## Acknowledgements

HepG2.117 cells were kindly provided by Prof. M. Nassal, University Hospital Freiburg im Breisgau, Germany. We would like to thank the following individuals for their expertise and assistance through various aspects of our study: Dennis Caluwé, Hongjun Chen, Jiadao Chen, Qianqian Chen, Heather Davis, Pascale Dehertogh, Guangyang Guo, Qinglin Han, Qiu Jin, Deborah Law, Qing Lu, Richard May, James Merson, Wendy Mostmans, Isabel Najera, Liping Shi, Karen Vergauwen, Gengyan Wang, Qun Wu, Guang Yang.

---

**Supplementary Figure 1.**
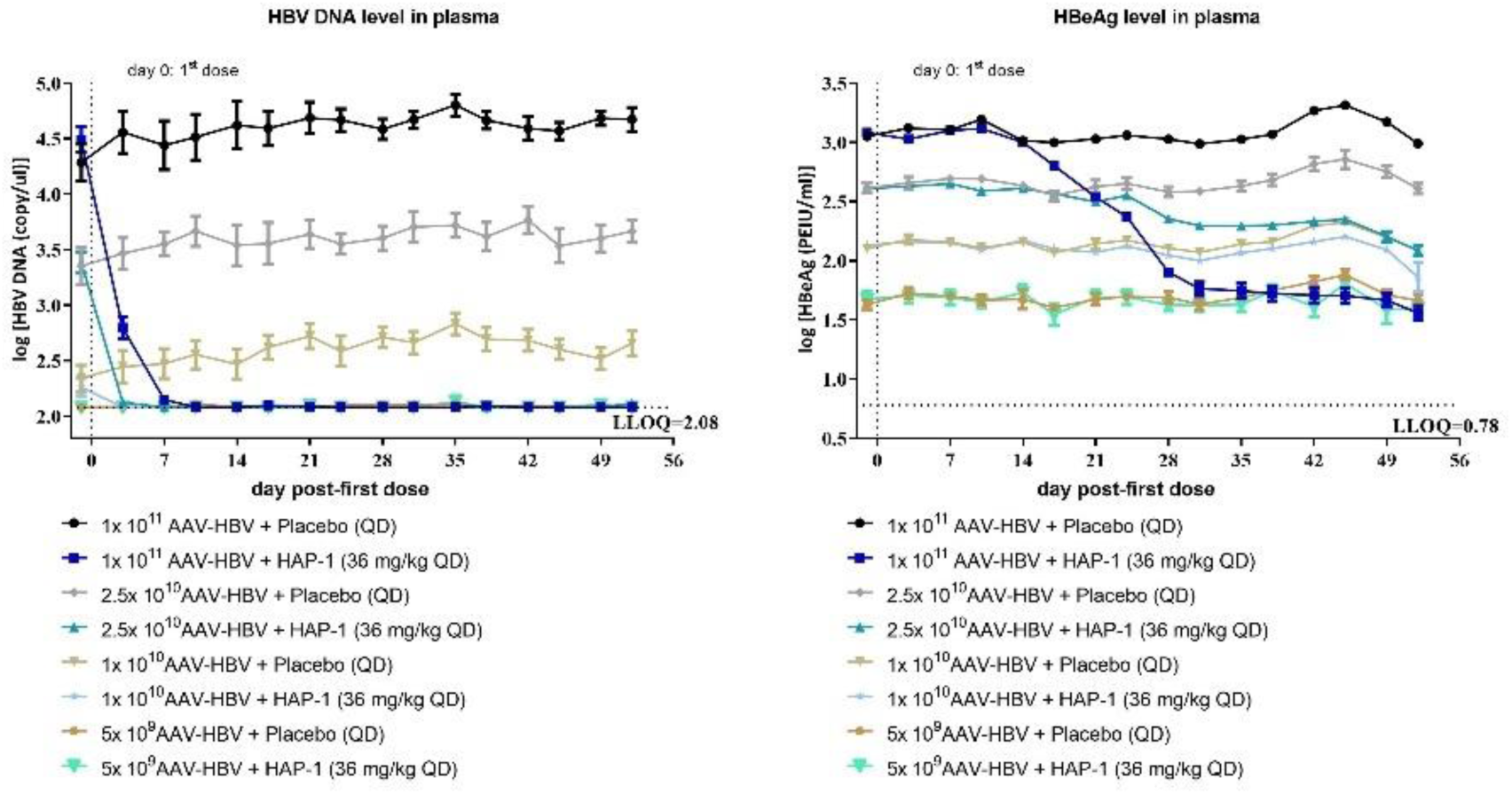
HBV DNA and HBeAg levels in mice treated with different doses of AAV-HBV. HBV DNA, HBeAg and ALT levels of C57BL/6 wt mice that were transduced with four different amounts of AAV-HBV (5x10^9^, 1x10^10^, 2.5x10^10^ and 1x 10^11^ AAV-HBV vg equivalents) treated orally for 56 days once daily (QD) with 36 mg/kg HAP-1 or placebo. QD once daily; wt wildtype; vg viral genomes; LLOQ lower limit of quantification. Data are shown as mean ± SEM and represent groups of 5 mice for the placebo treated control groups and 10 mice for HAP-1 treated groups.

**Supplementary Figure 2.**
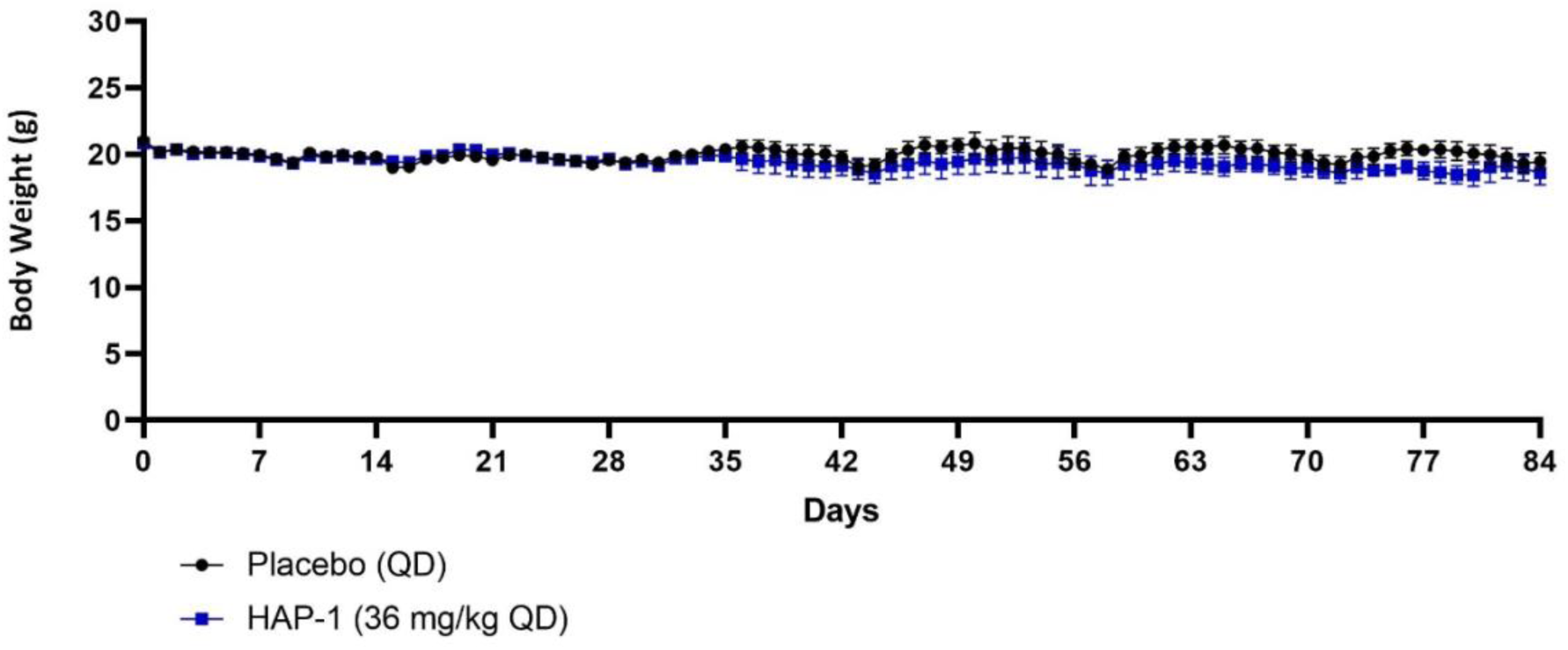
Body weight of HBV-infected chimeric mice treated with HAP-1 or placebo. HBV infected chimeric humanized liver mice were treated orally for 84 days once daily (QD) with 36 mg/kg HAP-1 or placebo. Body weight was monitored during treatment duration. Data are shown as mean ± SEM and represent groups of 3 mice for the placebo treated control groups and 4 mice for the HAP-1 treated groups.

**Supplementary data table 1:** limmaoutput_alltimepoints_paper (txt.file) available upon publication

