## SupplementaryMaterials for "Class A capsid assembly modulator apoptotic elimination of hepatocytes with high HBV core antigen level in vivo is dependent on de novo core protein translation"

[b]Infectious Diseases Discovery, Janssen Research and Development, 6F, Building 1 (North), Jinchuang Mansion, 4560 Jinke Road, Pudong, Shanghai, P.R. China, 201210

[c]Infectious Diseases Biomarkers, Infectious Diseases and Vaccines, Janssen Research and Development, Turnhoutseweg 30, 2340 Beerse, Belgium

[d]Infectious Diseases Discovery, Janssen Research and Development, 1600 Sierra Point Parkway, Brisbane, USA

* Current Address: Acerta Pharma B.V., A Member of the AstraZeneca Group, Pivot Park, Chesbrough (RK) Building, Kloosterstraat 9, 5349 AB, Oss, The Netherlands

[#] Current Address: Department of Microbiology and Immunology, McGill University

[†] Current Address: Retina Discovery, Cardiovascular & Metabolism Therapeutic Area, Janssen Research and Development, 1600 Sierra Point Parkway, 94005 Brisbane, USA

[¥] Current Address: AstriVax NV, Ambachtenlaan 1, B-3000 Leuven, Belgium

**SUPPLEMENTARY DATA**

**MATERIALS AND METHODS**

**Compounds**

HAP-1,2 and 3 were synthesized at WuXi AppTech, Shanghai, Peoples Republic of China, with a purity of >99%. GLS-4 was used as the positive high control for the induction of aberrant core structures and was synthesized in house. The GalNac-conjugated HBV siRNA was synthetized in a CRO. All compounds were prepared for research purposes only.

**Antiviral assay with HepG2.117 cells and cytotoxicity assay**

Details of the antiviral and cytotoxicity (HepG2) assays have been described previously (1). (Berke et al, 2017a). In brief, compounds at several concentrations, were added to cultured HepG2.117 cells (kindly provided by Dr. M. Nassal, University Hospital Freiburg, Germany). On day 4, HBV DNA was extracted from the cells and quantified by a quantitative polymerase chain reaction (qPCR) assay (1). Percent inhibition (50% effective concentration [$EC_{50}$] and 90% effective concentration [$EC_{90}$] values) was calculated using the difference in threshold cycle number between treated and control wells. Cultured HepG2 cells were incubated for 4 days with several concentrations of HAP1, 2 and 3. Cytotoxicity was evaluated by resazurin read-out, and 50% cytotoxic concentration ($CC_{50}$) and 90% cytotoxic concentration ($CC_{90}$) were calculated. Selectivity index was defined as the ratio of the mean $CC_{50}$ or $CC_{90}$ value of HAP1,2 and 3 in HepG2 cells to the mean $EC_{50}$ value of HAP1,2 and 3 assessed in HepG2.117 cells.

**High-content imaging HBV core protein aggregation assay**

Details of the high-content imaging core protein aggregation assay have been described previously (2). In brief, compounds were tested in dose range in presence of 1% dimethyl sulfoxide (DMSO) using a 25 µM or 100 µM start concentration and a serial dilution factor of ¼ in core expressing HepG2.117 cells. Compound plates with seeded cells were incubated for 72 hours at 37°C and 5% $CO_2$. After 72 hours, cells were fixated, permeabilized and core protein was detected with primary monoclonal mouse core antibody (Abcam Ab8637). After overnight incubation at 4°C, primary antibody was detected with Alexa Fluor-488 goat ant-mouse secondary antibody. Nuclei and cytoplasm were co-stained using Hoechst 33258 (nuclei) and HCS CellMask™ Deep Red (entire cells) solution. Images were captured via high content imaging using the Opera Phenix™ high-content screening system (Perkin Elmer), whereby image analysis via an Acapella script identified nuclei based on the Hoechst 33258 channel, cytoplasm segmentation was based on the HCS CellMask™ Deep Red channel and The texture feature was normalized to “% effect” such that the average over low control wells (i.e. non-compound treated core-expressing HepG2.117 cells) corresponds to 0% effect and the average over the high controls (i.e. 1 µM GLS-4 treated core-expressing HepG2.117 cells) corresponds to 100% effect. Non-linear log-logistic curve fitting was chosen for calculation of 50% effective concentration ($EC_{50}$) values using GraphPad Prism or Phaedra software which is an open source platform for data capture and analysis of high-content screening data (3).

**Core dependent apoptosis induction study in HepG2.117 cells**

After passaging, HepG2.117 cells were resuspended either in core expressing medium, containing DMEM supplemented with 2% Tet-system-approved fetal calf serum (FCS; Clontech), 2 mM alanyl-glutamine and 1x non-essential amino acids (NEAA; Sigma-Aldrich) or either in core suppressing medium, containing DMEM supplemented with 2% Tet-system-approved FCS, 2 mM alanyl-glutamine, 1x NEAA and 100 ng/ml doxycycline. 2,000 cells per well were seeded in compound-containing 384-well black poly-D-lysine PhenoPlate tissue culture treated plates with an optically clear bottom (Perkin Elmer).

HAP-1 and HAP-2 were tested in dose range in presence of 1.5% DMSO using 50 µM start concentration and a serial dilution factor of 1/3 in core expressing HepG2.117 cells or core suppressed HepG2.117 cells. Compound plates with seeded cells were incubated at 37°C and 5% $CO_2$ with media and compound refreshes performed every 3 to 4 days. Plates were incubated until day 28 with intermediate end points at day 14 and day 21 for parallel assay plates. Caspase 3/7 activity was measured via the CellEvent Caspase-3/7 Green detection reagent (Thermo Fisher Scientific). The CellEvent Caspase-3/7 green detection reagent was added to the cells in a final dilution of 1/400 and incubated for 1 hour at 37°C without $CO_2$. After incubation, images were captured using the CellVoyager CV8000 (Yokogawa) whereby image analysis using a custom Acapella (Perkin Elmer) script calculated the green fluorescent signal of the CellEvent Caspase-3/7 green detection reagent. Cell Viability was measured on parallel plates stained according to the high-content imaging HBV core protein aggregation assay using the cell count feature as a measure for cytotoxicity.

**Studies in AAV-HBV transduced and HBV-infected chimeric mice with humanized liver**

Male C57BL/6 and C57BL/6 SCID mice (age 5 weeks upon arrival) were injected with AAV-HBV virus diluted in PBS to $1.0\times10^{11}$ viral genome in 200 µL volume/mouse via tail veins. To study core protein dependency, mice were transduced with four different doses of AAV-HBV ($5x10^9$, $1x10^{10}$, $2.5x10^{10}$ and $1x10^{11}$ AAV-HBV vg equivalents). A vein blood sample was taken regularly (generally once per week) prior to treatment to evaluate HBV transduction efficiency by measurement of plasma HBV DNA, HBsAg, HBeAg, HBcrAg and ALT levels. Mice were randomly allocated into groups according to the plasma HBV DNA, HBsAg,

HBeAg level and body weights to achieve balanced distribution across groups. The animal welfare and euthanasia were conducted according to the standard operating procedures approved by Institutional Animal Care and Use Committee (IACUC).

**Quantification of HBV DNA in plasma of AAV-HBV transduced mice**

HBV DNA in mouse plasma was isolated with the QIAamp 96 DNA Blood Kit according to manufacturer's instructions (Qiagen, 51162). The isolated HBV DNA was quantified by qPCR using FastStart Universal Probe Master (Roche, 04914058001). The standard curve DNA of $10^7$ copies/μL was prepared by 60-fold dilution of 5 ng/μL pAAV2-HBV1.3 plasmid DNA, followed by a 10-fold serial dilution in AE buffer to generate standards ranging from $10^7$ to 10 copies/μL. HBV DNA levels were interpolated according to the standard curve. The LLOQ of the HBV DNA assay is determined as 120 copies/μL plasma.

**HBsAg and HBeAg measurement in plasma of AAV-HBV transduced mice**

For the detection of HBsAg, mouse plasma samples were first diluted 50-fold, then further diluted another 24-fold. Fifty μL of the 1200-fold diluted plasma was used for determination of HBsAg levels using the HBsAg ELISA kit (Autobio, CL 0310) following the supplier's manual.

For the detection of HBeAg, mouse plasma samples were diluted for 50-fold (4 μL plasma + 196 μL PBS). Fifty μL of the 50-fold diluted plasma was used for determination of HBeAg using the HBeAg ELISA kit (Autobio, CL 0312) following the supplier's manual.

**Measurement of HBcrAg in plasma of AAV-HBV transduced and chimeric mice**

Quantitative levels of HBV core-related antigen (HBcrAg) were determined using the Lumipulse G HBcrAg assay, which measures simultaneously denatured HBeAg, HBcAg and the precore protein p22cr (aa −28 to aa 150). Samples were handled according to the manufacturer's instructions. HBcrAg levels are quantified in U/mL.

**ALT activity assay in plasma of AAV-HBV transduced mice**

ALT levels in mouse plasma were detected using the ALT Activity Assay Kit (Sigma, MAK052) following the supplier's manual.

**Liver Immunohistochemistry with liver samples from AAV-HBV transduced mice**

Formalin-fixed, paraffin-embedded liver samples were cut and labelled for immunohistochemistry (IHC) with a polyclonal rabbit anti-HBcAg antibody (Abcam, ab115992, dilution factor 1:250) or a polyclonal horse anti-HBsAg antibody (Abcam, ab9193, dilution factor 1:250). Isotypes-matched antibodies were used as control primary antibodies and HBV-negative liver tissue was used as control tissue. Positive staining was observed under a microscope and representative photos were captured under 200-folds view.

**Liver Immunofluorescence with liver samples from AAV-HBV transduced mice**

Formalin-fixed, paraffin-embedded liver samples were sectioned at 5µm, mounted on SuperFrost Plus glass slides (VWR, 631-9483) and immunofluorescently stained using a VENTANA Discovery Ultra (Roche) for HBsAg/Ki67/DAPI. Alternatively, TUNEL assay (Invitrogen, C10617) was performed manually following the vendor guidelines. All immunofluorescent stained sections were covered with mounting media (DAKO, S3023) and scanned on a NanoZoomer S60 (Hamamatsu). Isotypes-matched antibodies were used as control for primary antibody and HBV-negative liver tissue was used as control tissue and for biomarker positivity interpretation.

| | **SUPPLIER** | **REFERENCE** | **CLONE** | **DILUTION** |
|---|---|---|---|---|
| **HBsAg** | Abcam | Ab859 | 3E7 | 1:1 |
| **Ki67** | Abcam | Ab15580 | - | 1:500 |

**Liver immunofluorescence cell segmentation and quantification.**

Immunofluorescent images were subjected to cell segmentation using HALO image analyses software (Indica Labs) and CytoNuclear FL module to allow the identification of single cells and biomarker expression. Briefly, tissue boundaries and individual cells were identified by the presence of nuclear dye (DAPI) positivity. Afterwards, starting from the localization of a nucleus, the software gradually expands the area to identify the cell boundary (cytoplasm), where the biomarker intensity is calculated. Based on the overall intensity distribution or sample controls, cell positivity for HBsAg, TUNEL and Ki67 was determined to assign cellular phenotypes. Fraction of biomarker phenotypes was calculated over the total number of cells identified (number of DAPI positive nuclei in liver section).

**Cytokine detection**

Mouse plasma samples were used unprocessed, mouse liver samples were homogenized by bead beating on a TissueLyser twice during 3 min at 25 Hz/s after addition of a 5mm stainless steel bead (Qiagen, 69989) and 450 µl homogenate buffer (0.1% Triton X-100 and 2.5 mM EDTA in DPBS, supplemented with protease inhibitor (Roche, 05056489-001)). Homogenized samples were centrifuged for 15 min at 12000 x g after which supernatant is collected and normalized based on total protein content.

Detection of cytokine and chemokine biomarkers was done with Luminex xMAP bead-based assay platform, in combination with Milliplex kits (Merck-Millipore, MCD8MAG48K-PX15 and MCYTOMAG-70K) following the manufacturer's manual. In brief, samples are brought in contact with magnetic beads which are each coated with a specific capture antibody. After an analyte is captured by the beads, a biotinylated detection antibody is introduced and the reaction mixture is then incubated with Streptavidin-PE conjugate, the reporter molecule, to complete the reaction on the surface of each bead. Each individual bead is identified by its ‘bead signature’ and quantified based on fluorescent reporter signals using a Biorad Bio-Plex 200 system. Standard curves were generated with standards supplied with the kits, a best-fit line was determined by regression analysis using 4PL logistic curve-fit.

**Microarray**

Mouse liver (10 mg) was homogenized by bead beating on a TissueLyser twice during 60s at 20 Hz/s after addition of two 3.5mm stainless steel beads (Qiagen, 69989) and 450µl RLT Plus lysis buffer (Qiagen, 1053393). After homogenization, a total of 400µl lysate was used for RNA extraction on a QiaSymphony liquid handling system using the QIAsymphony RNA extraction kit (Qiagen, RNA_CT400), according to manufacturers’ instructions.

Transcriptome analysis was done using the Mouse Clariom S Assay (Thermo Fisher Scientific, 902930), in combination with the GeneChip WT PLUS Reagent kit (Thermo Fisher Scientific, 902280). A total of 100 ng RNA was used as input and downstream process was done according to the manufacturer’s guidelines. In brief, double-stranded cDNA is first synthesized using random primers, and then used as template for in vitro transcription (IVT). The cRNA is used as input for a second round of first strand cDNA synthesis, producing single stranded sense cDNA. After fragmentation and end-labeling, the targets are hybridized to a multi-sample Clariom S plate array on a GeneTitan Multi-Channel Instrument (Thermo Fisher Scientific).

Gene expression values were normalized by robust multiarray average normalization on the microarray probe-level data, and downstream analysis was done in R version 3.4.2.

Unsupervised analysis using spectral maps was performed to remove technical outliers. A supervised analysis was performed using the limma package (4) for comparisons between treatment groups across each timepoint and corrected p-values for multiple testing across genes $\leq 0.05$ were considered significant. The affected pathways were analyzed with MLP (mean log P analysis) with GO Biological Process and Ingenuity Pathway Analysis (Qiagen). The considered cutoffs for MLP were lower (5) and upper (250) threshold for gene set size where 7100 pathways from the Biological Process and 667 pathways from IPA were used.

**In vivo depletion of CD8+ T cells.**

Male AAV-HBV mice were intraperitoneally (i.p.) injected with anti-CD8β (BioXcell, 53.5.8) or IgG1 (BioXcell, HRPN) antibodies at 100 µg/mouse on day -4 and day -3 before compound treatment, followed with QW administration until day 39. Antibody injections were strengthened to Q2W (biweekly) from day 39 to day 56 to maintain efficient elimination of CD8+T cells. Concurrently AAV-HBV mice were orally administered with CAM-A (JNJ-3809, 20mg/kg, QD) compound from day 0 until day 56. CD8+ T cell frequency in CD3+ T cells were monitored in blood at weekly bases by flowcytometry using anti-mouse CD8α antibody (BD, 53.6.7).

**Studies in HBV-infected chimeric mice with humanized liver**

Male HBV-Genotype-C-infected uPA/SCID chimeric mice (PXB-Mouse®; 21 to 25 weeks of age; weight 17.8 to 23.0 g) were from PhoenixBio (Higashi-Hiroshima, Japan). All mice had blood human albumin levels above 7.8 mg/mL and serum HBV DNA levels above $10^6$ copies/mL. Median (range) HBV DNA levels across all randomized mice was 8.0 (6.6–8.8) $\log_{10}$ copies/mL. Mice received food and water *ad libitum*.

The use of the animals for this study was approved by the Animal Ethics Committee of PhoenixBio (Resolution No.: 1495). All the experimental procedures used to treat live animals in this study were approved by the Animal Ethics Committee of PhoenixBio.

**Determination of Blood Human Albumin**

At all blood samplings, human albumin levels were determined using the clinical chemistry analyzer (BioMajesty™ Series JCA-BM6050, JEOL) on saline-diluted blood, and latex agglutination immunonephelometry (LZ Test "Eiken" U-ALB, Eiken Chemical).

**Serum HBV DNA Quantification**

HBV DNA was extracted from serum using the SMITEST EX-R&D Nucleic Acid Extraction Kit (Medical & Biological Laboratories), dissolved in nuclease-free water, after which real-time polymerase chain reaction (PCR) was performed using the TaqMan™ Fast Advanced Master Mix (Applied Biosystems, Thermo Fisher Scientific) and ABI Prism® 7500 sequence detector system (Applied Biosystems).

**Serum HBsAg and HBeAg Quantification**

Serum HBsAg and HBeAg concentrations were determined by SRL (Tokyo, Japan) based on the ChemiLuminescence ImmunoAssay (CLIA) developed by Abbott (Abbott Park, IL; ARCHITECT® SYSTEM).

**Immunofluorescence stain for the detection of core and HBsAg in cryosections of mouse liver**

Fresh frozen liver samples were fixated for 10 minutes in PFA 4%. After a PBS wash, sections of 10 µm were cut, mounted on SuperFrost Plus glass slides (VWR, 631-9483) and immunofluorescently stained using a VENTANA Discovery Ultra (Roche) for HBsAg/DAPI or Core/DAPI. All immunofluorescent stained sections were covered with mounting media

(DAKO, S3023) and scanned on a NanoZoomer S60 (Hamamatsu). Isotypes-matched antibodies were used as control for primary antibody and HBV-negative liver tissue was used as control tissue and for biomarker positivity interpretation.

| | SUPPLIER | REFERENCE | CLONE | DILUTION |
|---|---|---|---|---|
| **HBsAg** | Abcam | Ab859 | 3E7 | 1:1 |
| **HBV Core Ag** | DAKO | B0586 | - | 1:2000 |
