## Supplementary material for "Class A capsid assembly modulator apoptotic elimination of hepatocytes with high HBV core antigen level in vivo is dependent on de novo core protein translation": Abbreviations

| | |
|---|---|
| AAV | Adeno-associated virus |
| ALT | Alanine Amino Transferase |
| CAM | Capsid assembly modulator |
| CAM-A | Capsid assembly modulator (inducing Aberrant structures) |
| CAM-E | Capsid assembly modulator (inducing Empty structures) |
| cccDNA | Covalently closed circular DNA |
| CD8 | cluster of differentiation 8 (transmembrane glycoprotein, co-receptor for T-cell receptor) |
| CD74 | cell surface receptor for cytokine MIF |
| CHB | chronic hepatitis B |
| DAPI | 4′,6-diamidino-2-phenylindole (a blue-fluorescent DNA stain) |
| $EC_{50}$ | 50% effective concentration |
| ELIspot | enzyme-linked immunosorbent spot |
| ER | endoplasmatic reticulum |
| ETV | Entecavir |
| FACS | Fluorescence-activated cell sorting |
| FAS | FS-7-associated surface antigen, a death receptor on the surface of cells |
| FASL | FAS Ligand, a key pro-apoptotic molecule |
| GalNac | N-Acetylgalactosamine, an amino sugar derivative of galactose |
| Granzyme B | one of the serine protease granzymes found in the granules of natural killer cells |
| H2-Ab1 | histocompatibility 2, class II antigen A |
| H2-Eb1 | histocompatibility 2, class II antigen E beta |
| H2-Q7 | histocompatibility 2, Q region locus 7 |
| hAlb | human Albumin |
| hALT | human ALT |
| HAP | heteroaryldihydropyrimidine |
| HBV | hepatitis B virus |
| HBcrAg | Hepatitis B core-related antigen |
| HBeAg | Hepatitis B e antigen |
| HBsAg | Hepatitis B surface antigen |
| IFN-γ | Interferon gamma, cytokine crucial in inflammatory response |
| IP10 | interferon-inducible protein 10 (or CXCL10) is a chemokine of the CXC family |
| IU | international units |
| Ki67 | nuclear protein associated with cellular proliferation |
| LLOQ | Lower limit of quantification |
| MHC-1 | major histocompatibility complex class I molecules |
| MIF | macrophage migration inhibitory factor |
| MIG | Monokine induced gamma interferon (cytokine that belongs to CXC chemokine family) |
| MOA | Mechanism of action |
| NA / NUC | Nucleos(t)ide analogue |
| p-value | probability, measures likelihood of observed difference between groups is due to chance |
| pgRNA | Pregenomic RNA |
| pol-pgRNA | Polymerase-bound pgRNA |
| QD | Once daily |
| RANTES | regulated upon activation, normal T cell expressed and secreted (alias CCL5), a potent chemoattractant cytokine |
| rcDNA | Relaxed circular DNA |
| s.c. | subcutaneous |
| SCID | severe combined immunodeficiency disease |

| | |
|---|---|
| siRNA | small interfering RNA |
| TDF | tenofovir disoproxil |
| TUNEL | Terminale deoxynucleotidyltransferase dUTP nick-end labeling |
| UPR | Unfolded Protein Response |
| vg | viral genomes |
| WHO | World Health Organisation |
| wt | wild type |
